# Breaking Symmetry in Walking: Drosophila Recruits Task- and Leg-Specific Motor Modules for Curve Walking

**DOI:** 10.64898/2026.10.22.760936

**Authors:** E. A. Gorostiza, R.D. Custódio, M. Dübbert, Kei Ito, T. Bockemühl, A. Büschges

**Affiliations:** Institute of Zoology, Faculty of Mathematics and Natural Sciences, University of Cologne, Cologne, Germany

**Author notes:** co-senior author (T. Bockemühl, A. Büschges).

**Keywords:** Walking, Kinematics, Motor control, Steering, *Drosophila melanogaster*

## Abstract

Walking animals must continuously reorient to navigate their environment, yet how leg kinematics are reconfigured to produce directional changes of varying magnitude remains poorly understood. Here, we used high-resolution leg tracking to characterize steering behavior in freely walking *Drosophila*. We focus our analysis on two major strategies, arcs and turns. Although arc and turn trajectories are statistically separable, several parameters suggest an underlying spatial continuum spanning both behavioral categories, involving a change in the steering mechanism rather than a discontinuity in the kinematic space. Arcs, representing shallow curves, and were generated primarily through: (1) graded shifts in the posterior extreme position (PEPs) of the front and inside middle legs, and bilateral asymmetries in stance amplitude, lengthening on the outside (2) and shortening on the inside (3) while step timing and ipsilateral coordination remained at straight-walking baseline. Turns, in contrast, were associated with higher steering demands and relied on: (4) progressive displacement of the anterior extreme positions (AEPs) in front and middle legs, (5) extreme reduction of inside middle and hind leg amplitudes to anchor the turning pivot, (6) skipped steps of inside middle and hind legs, (7) lateral steps of inside middle legs, and (8) spontaneous appearance of synchronized backward-stepping in particular legs beyond a critical turning threshold. Finally, a common kinematic configuration emerges at maximal turning demand. These results suggest a hierarchical, bipartite organization of steering: PEP- and AEP-driven mechanisms operating across moderate and severe trajectory curvatures, and a shared saturated motor state at the behavioral extreme that links descending steering commands to segment-specific leg kinematics in a freely walking insect

**Graphical abstract:** 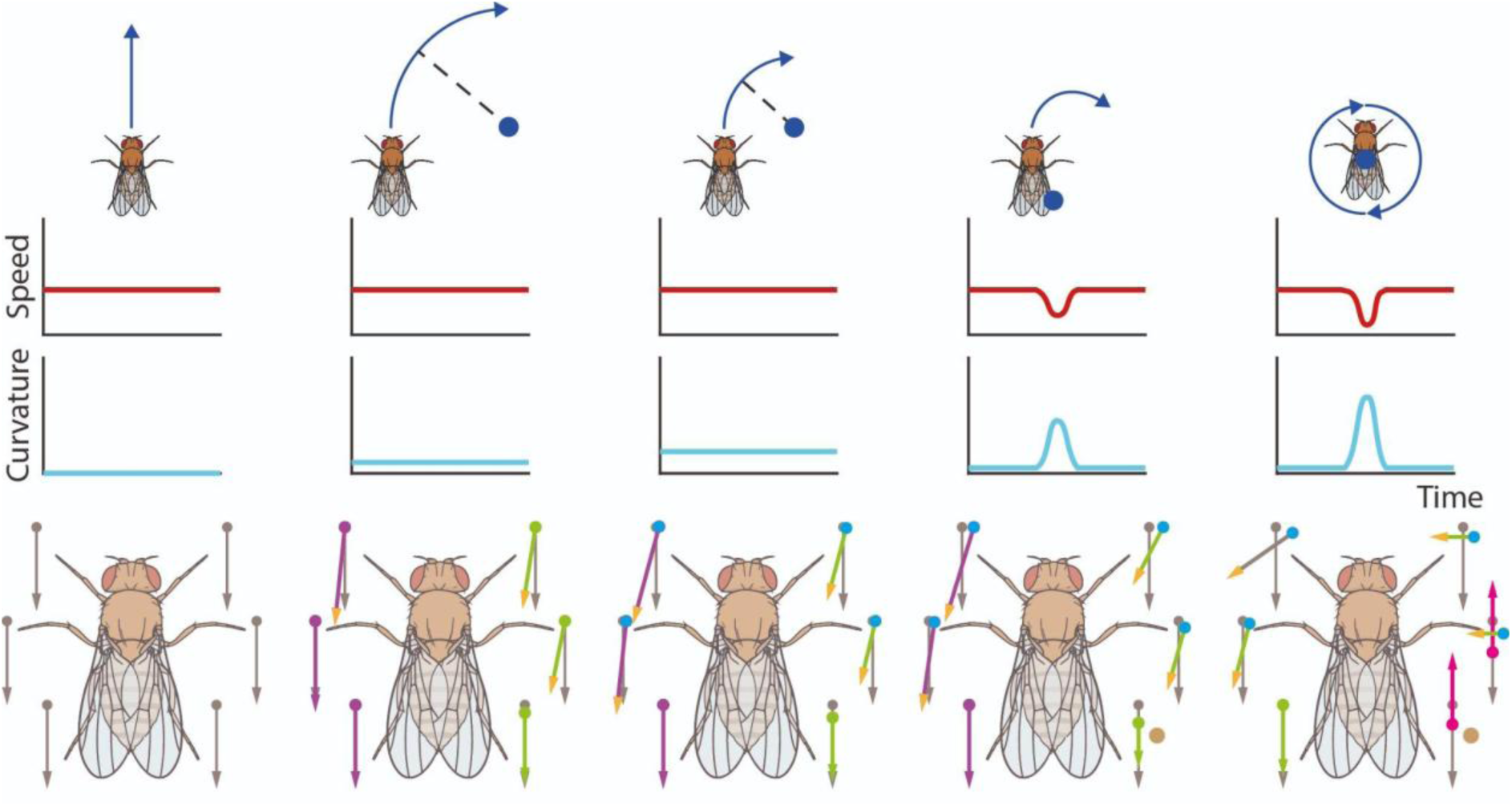

**Highlights:**

- *Drosophila* flies use several strategies to change direction, including **arcs** and **turns**
- **Arcs** have constant curvature, speed and constant angular velocity, while **turns** require a decrease in speed before changing direction and their angular velocity tends to be higher.
- We identified eight leg kinematic mechanisms that contribute to these strategies.
- Some kinematic parameters vary smoothly and in a continuous fashion between mechanisms.

## Introduction

Straight walking has provided a tractable paradigm to understand basic principles of interleg coordination. However, it constitutes a special case; animals rarely move in perfectly straight lines. Navigating environments, tracking odors, or pursuing visual targets all demand frequent and often abrupt reorientations. Understanding how nervous systems reconfigure leg movements during walking to change heading, while preserving gait stability and propulsion remains a central question. Changing heading is fundamental for all walking, regardless of the number of legs. For curved locomotion, bipeds, quadrupeds, hexapods and myriapods produce breaks in left–right symmetry. The way this asymmetry is implemented differs with posture, number of legs or body flexibility (Courtine and Schieppati 2003; Molkov et al. 2026; Jindrich and Full 1999; Aoi et al. 2016; Walter 2003; Dürr and Ebeling 2005). In centipede-like systems, for instance, available evidence emphasizes body-axis flexibility, gait transitions, and strong local body-leg coordination, suggesting that steering in these elongated multi-legged animals may depend more on embodied mechanics than in walkers with few legs (Aoi et al. 2023; Mingchinda et al. 2023; Dionne et al. 2026).

Turning behavior has been characterized kinematically in several insects, including honeybees, cockroaches, beetles, and stick insects (Zolotov et al. 1975; Franklin et al. 1981; Frantsevich and Mokrushov 1980; Gruhn et al. 2009). These studies identified two overlapping mechanisms: bilateral asymmetric modulation of step amplitude and frequency, as well as the transient fixation of an inside leg as pivot point. Detailed analyses in cockroaches and stick insects further showed that outside legs tend to push the body forward through the turn while inside legs are repositioned to pull it toward the new heading (Jindrich and Full 1999), with leg-specific changes in anterior and posterior extreme positions, stance and swing timing, and interleg phase relationships (Jindrich and Full 1999; Dürr and Ebeling 2005; Gruhn et al. 2009, 2016). However, most of this work relied on tethered or restrained preparations and on a limited number of turning categories, leaving open how these low-level adjustments are organized across the full range of natural steering maneuvers.

Building on a long history of kinematic research in *Drosophila*, from classical observational studies (Götz and Wenking 1973; Strauss and Heisenberg 1990) to high-resolution quantitative analyses (Mendes et al. 2013; Wosnitza et al. 2013) the fruit fly provides an established model for dissecting the motor control of hexapedal walking and revisiting these questions. Forward walking in *Drosophila* is characterized by a speed-dependent continuum of interleg coordination without segregating into traditional discrete gaits. Instead, it forms a continuous manifold, a low-dimensional geometric space where all possible coordination patterns are parameterized along a primary axis defined by the speed-dependent modulation of stance duration (DeAngelis et al. 2019; Wosnitza et al. 2013; Szczecinski et al. 2018). Beyond straight locomotion, directional changes during freely walking could be rapidly changed by saccade-like body turns (Geurten et al. 2014) or steering maneuvers that implement asymmetric modulations of the body size. Recent work has identified descending neurons related to these maneuvers, showing that activation of specific cell types drives coordinated shortening of strides on the inside of a turn or lengthening on the outside, scaling with turning angular velocity (Yang et al. 2024). These circuit- and kinematic-level findings imply that steering in *Drosophila* can be tuned continuously, yet it remains unclear whether the diversity of curved walking behavior reflects this same single underlying kinematic continuum or several kinematically distinct motor programs, and how leg-specific spatial and temporal parameters are reorganized across this repertoire.

Here, we combined high resolution videography of freely-walking flies with leg tracking (Mathis et al. 2018) to systematically characterize curved walking in *Drosophila*. Consistent with established insect maneuvers, we identified three distinct steering strategies that map closely to the swerves, pivots, and body saccades described in previous reports. Here, we focus on two of these modes, arcs and turns, to dissect how local leg kinematics are reconfigured as a function of angular velocity and turning angle. Our results revealed both continuously graded and at least four categorically distinct leg-level mechanisms underlying steering, providing a kinematic framework to link descending motor commands to the diversity of turning behavior in a freely walking insect

## Results

### Flies predominantly use three strategies to generate changes in direction

To better understand curve walking behavior in *Drosophila*, we captured videos of free-walking flies at high spatio-temporal resolution. In curve walking sequences, we identified three main strategies we called adjustments, arcs, and turns (Fig1 A-B). Adjustments were characterized by fast, but small, directional changes, without clear deceleration of translational speed, flanked by straight walking; these resembled walking saccades (Geurten et al. 2014). They comprised around 150 milliseconds, and were produced within one step of the hind leg inside of the curve. We decided to not further investigate them, and focused on arcs and turns. These two categories resembled in principle those already acknowledged in flies (Yang et al. 2024; Strauss and Heisenberg 1990; Cruz et al. 2021) and other insects (Zolotov et al. 1975; Franklin et al. 1981). However, our classification relies operationally on trajectory geometry and global dynamics. By framing steering manoeuvres along a natural geometric continuum of curvature, from broad arcs to sharp turns, we could evaluate whether this spatial progression reflects a single continuous motor program or distinct kinematic strategies. An arc was defined as a walking sequence with constant speed, curvature and angular velocity (Fig1 C-D) creating a shallow curve with the pivot point far from the body (Yang et al. 2024). In contrast, a turn was characterized by a decrease in translational speed during which a sudden increment in curvature and angular velocity happened (Fig1 D) (Cruz et al. 2021; Yang et al. 2024; Strauss and Heisenberg 1990). Turns could reach higher angular velocities and cover usually less distance than arcs (Fig1 C). Also, the decrease in speed typically preceded the peak of angular velocity (Fig1 D). Due to these changes in speed, speed-matching was not possible between turns and straight walking in further analyses, as we did for arcs.

We decided to explore low level kinematics of arcs and turns. After cropping the videos, tarsal tip positions were determined using DeepLabCut, (DLC,(Mathis et al. 2018). For each step, we extracted anterior and posterior extreme positions (AEPs and PEPs, respectively -see Videos 1-5-), stance amplitude, stance, swing, and step durations, as well as phase relationships between adjacent legs and compared these parameters to straight walking. As in previous *Drosophila* studies (Yang et al. 2024), we found changes in several parameters (Fig1 E-F). During arcs, especially both front legs, inside and outside of the curve (IFL and OFL, respectively), and the inside middle leg (IML) rotated the orientation of their stance trajectories into the direction of the curve, while stance orientation in the remaining legs did not change appreciably. Interestingly, AEPs and PEPs had a spatial dispersion comparable to straight walking (Fig 1E, red boxes). In all legs, stance amplitudes were higher in the outside legs than in the inside ones. For the outside middle leg (OML) and outside HL (OHL) stance amplitudes increased, while the amplitudes on the corresponding inside legs decreased. However, the stance amplitudes of outside FL (OFL) appears unaffected, while inside decreased (Fig 1F, red boxes). Swing duration showed only clear difference for the ML, being longer for the OML and shorter for the IML (Fig 1F, red boxes). These differences were compensated by stance duration, in which the opposite effect was observed, keeping the overall step duration on the level of straight walking. A subtle trend emerged in the phase relationships between inside and outside legs (Fig1 G, left plot, red boxes). In comparison to straight walking, the OFL was delayed relative to the IFL, whereas the ML and HL exhibited the opposite relationship. Nevertheless, flies appeared to maintain a high degree of coordination stability, comparable to straight walking, as these phase shifts remain minimal, with median shifts staying close to zero and showing low variability during arcs (Fig1 G). Ipsilateral phase relationships during arcs remained unchanged compared to straight walking (Fig1 G, right plots) indicating that the anteriorly-directed swing phase progression along each side of the body is largely preserved (Szczecinski et al. 2018). This suggests that during large-radius curves, turning dynamics are handled primarily through bilateral adjustments rather than ipsilateral reconfiguration.

**Figure 1.**
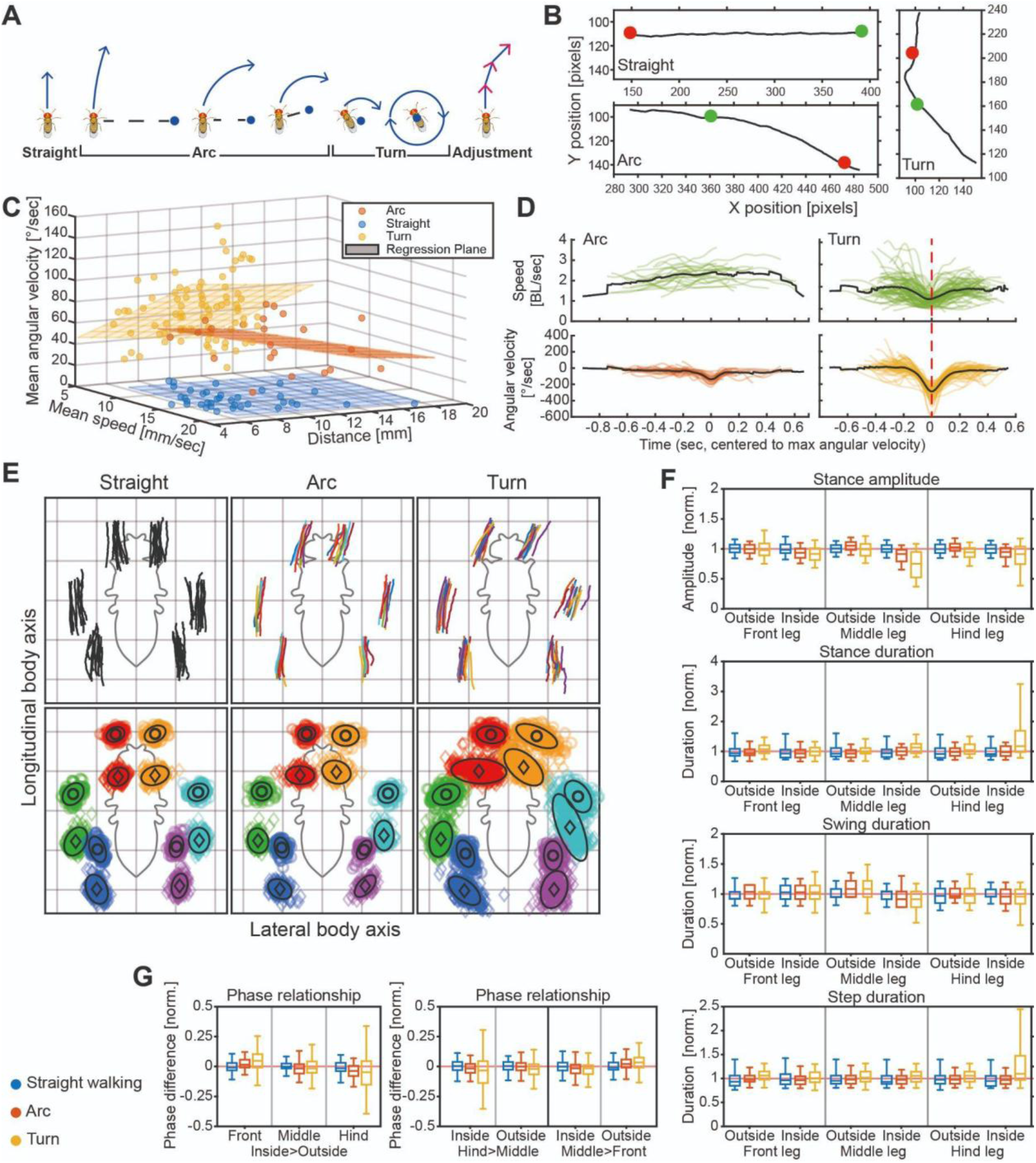
Kinematic characterization of arcs and turns. **A)** Schematic of the different main strategies for changing directions found in our video samples. **B)** Examples of trajectories selected for the analysis. On the top left, Straight walking; bottom left, Arc; On the right, Turn. Green and red dot, mean the start and end points, respectively, of the selected sequences. **C)** Average angular velocity, distance, and average translational speed during the sequences. **D)** Translational speed and angular velocity along each trajectory analysed in this section, centered to the maximum angular velocity point on it. For visualization and averaging, traces were Gaussian-smoothed over a temporal window equal to 30% of the sequence length. **E)** Average stance trajectories (top) and distribution of AEP (circles) and PEP (diamonds) of all flies walking straight (N=11, n=37), performing arcs (N=22) or turns (N=89) in a fly-centered coordinate system. Black circles and diamonds represent mean AEP and PEP positions, and the black ellipses are the 95% percentiles. **F)** Kinematic parameters calculated for complete steps for straight walking, arc and turns. All data shown here have been normalized to the mean values in the straight walking condition (indicated as 1, light red reference line). Box plots (whiskers indicate 1% and 99% percentiles) are combined normalized data for all flies. **G)** Phase relationships between ipsilateral (Hind>Middle and Middle>Front) and contralateral leg pairs (Front, Middle, Hind, Inside>Outside), Inside and Outside, refers to the legs on the inside or outside of the curvature trajectory.

In contrast to the stability observed during arcs, turns triggered a significant amplification of these kinematic shifts accompanied by a marked increase in overall variability across all evaluated parameters (Fig 1E-G, yellow boxes). These changes were particularly evident in the inside HL (IHL), which often acted as a pivot point for the manoeuvres. We also observed that IHL may skip a step or perform an extremely short one, a behavior also observed in the IML. This aligns with classic principal turning models for insects, which propose either an increase in step amplitude in the outside legs or the fixation of an inside leg to act as a pivot (Zolotov et al. 1975; Franklin et al. 1981; Frantsevich and Mokrushov 1980). Here, this was quantitatively supported by a sharp decrease in stance amplitude for both the IML and IHL, coupled with a substantial increase in HL stance duration (Fig 1F). Consequently, data dispersion was significantly higher than in straight walking and arcs (Fig1 G, yellow boxes). It is worth mentioning that the turn sequences analyzed here included a step before and after the turn itself, which might contribute to the observed variability and dampen the changes observed

To explore whether these strategies stem from an underlying kinematic continuum like coordination patterns in straight walking, we combined an optomotor stimulus (Yang et al. 2024; Dürr and Ebeling 2005; Gruhn et al. 2016, 2009) with a free-walking arena to enrich the dataset with curved trajectories. Video-recording flies under various stimulus rotational speeds yielded over 1100 sequences of straight walking, arcs, and turns. Kinematic analysis confirmed that turns exhibit lower mean speeds, higher mean angular velocities, and shorter distances than arcs (Fig. S1A, top row; Fig. S1B). These traits are accompanied by distinct trajectory geometries (SD curvature AUC = 0.995) and translational dynamics (mean speed AUC = 0.924), which differentiate the two strategies more strongly than rotational magnitude (mean angular velocity AUC = 0.741).

### Stance trajectory shift in arcs depends primarily on PEP shift

We then tracked for each sequence tarsal tips positions, and used angular velocity of the fly to sort them out, spanning for the arcs from 40°s^−1^ to 220°s^−1^. As expected from previous works (Yang et al. 2024), when compared to straight walking, increments in angular velocity correlated with increments in stance trajectory shifts. However, on average, these shifts occurred primarily in both FL and the IML, whereas the OML shifted minimally until angular velocities exceeded 100°s^−1^ (Fig 2A, Fig S2A). Interestingly, these adjustments were driven by clear PEP displacements as angular velocity increased, while the AEP showed only subtle changes. This was evident in the spatial dispersion profiles. While the AEP remained as tightly clustered as in straight walking, the PEP for both FL and the IML dispersed progressively with higher angular velocities (Fig 2A, S2A). In this regard, spatial adjustments varied across leg pairs (Fig 2B, S2A, S3 top). In the FL, PEP showed substantial and consistent displacement, outward-posteriorly on the outside and outward-anteriorly on the inside, whereas on both AEP only minor shifts could be detected. For the ML and HL, lateral symmetry broke down. The IML showed a constantly anteriorly shifted PEP, but the AEP minimally moved from baseline. The OML changed subtly, moving its PEP outward only past 160°s^−1^. The OHL remained unaltered, but the IHL shifted along the anteroposterior axis. These coordinated shifts systematically alter stance trajectories and their amplitudes; OFL and OML trajectories lengthened, whereas IML and IHL trajectories shortened. A similar PEP shift occurs in stick insects, where the transition to curved walking alters the PEP during the first step of both FL before affecting the AEP (Dürr and Ebeling 2005).

**Figure 2.**
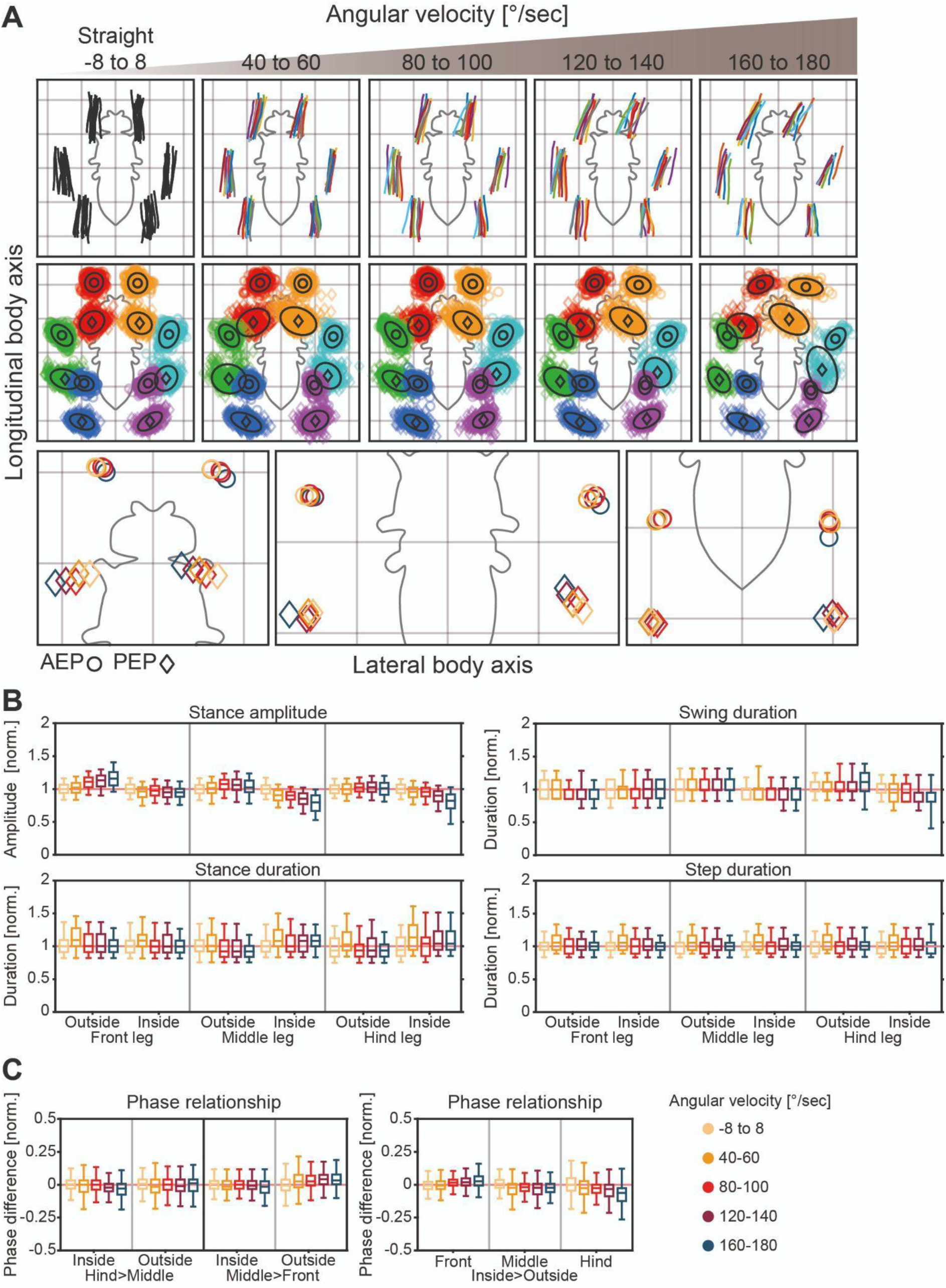
Low kinematic analysis of arcs at different angular velocities. **A)** Average stance trajectories (top) and distribution of AEP (circles) and PEP (diamonds) of all flies walking straight (-8° to 8°, N=11, n=120), or performing arcs (N=11, n=527) in a fly-centered coordinate system. **B)** Kinematic parameters calculated for complete steps for straight walking, and arc. All data shown here have been normalized to the mean values in the straight walking condition (indicated as 1, light red reference line). Box plots (whiskers indicate 1% and 99% percentiles) are combined normalized data for all flies. **C)** Phase relationships between ipsilateral (Hind>Middle and Middle>Front) and contralateral leg pairs (Front, Middle, Hind, Inside>Outside), Inside and Outside, refers to the legs on the inside or outside of the curvature trajectory. Only a selection of the total range of explored angular velocities is shown here, see supplementary figure 1 and 2 for more details. N and n, are from the complete experiment.

Step duration remained stable across angular velocities due to compensatory changes in swing and stance phases, reflecting robust temporal control (Fig 2C, Fig S3 middle). Although values were comparable to straight walking, the ML and HL showed distinct contralateral trends: outside legs exhibited longer swings and shorter stances, while inside legs displayed the inverse pattern .

Bilateral phase analysis (Fig. 2C, S3 bottom) revealed that median values shifted progressively from the zero-baseline as angular velocity increased in a pair-specific manner. In the FL, a positive shift indicates a velocity-dependent delay of the outside leg relative to the inside one, while the HL exhibited a progressively stronger negative displacement. Although the ML also displayed a negative phase shift, this displacement remained relatively constant and decoupled from changes in angular velocity.

Ipsilateral phase analysis during arcs revealed pair-specific modulations rather than a rigid global command (Fig. 2C, S3 bottom). While outside HL>ML and inside ML>FL couplings remained stable, probably to preserve propulsive flow, other pairings shifted with angular velocity. Specifically, the outside ML>FL pairing showed a consistent positive deviation, delaying the front leg likely to accommodate the larger external trajectory. Conversely, the inside HL>ML coupling displayed a significant negative shift at high angular velocities (>120°s^−1^), likely modulating hind-leg timing to facilitate rotation. These results indicate that the fly selectively tunes specific ipsilateral phases to meet the mechanical demands of increasing curvature while preserving forward progression.

### During turns, stance trajectory shift depends primarily on AEP shift

Kinematic analysis of turn sequences from the same dataset revealed a diverse range of sub-strategies, including skipped and backward steps, particularly in the IML and IHL. Because of their distinct kinematic profile, turns containing backward steps were excluded to focus solely on maneuvers maintaining forward progression across all legs. Unlike arcs, turns lacked sufficient data between 40°s^−1^ and 60°s^−1^; thus, our analysis spans from 60°s^−1^ to 220°s^−1^. Consistent with previous work (Yang et al. 2024) and our present findings, increasing angular velocity correlated with stance trajectory shifts, which were markedly larger during turns than arcs (Fig. 3A, Fig. S2 middle).

**Figure 3.**
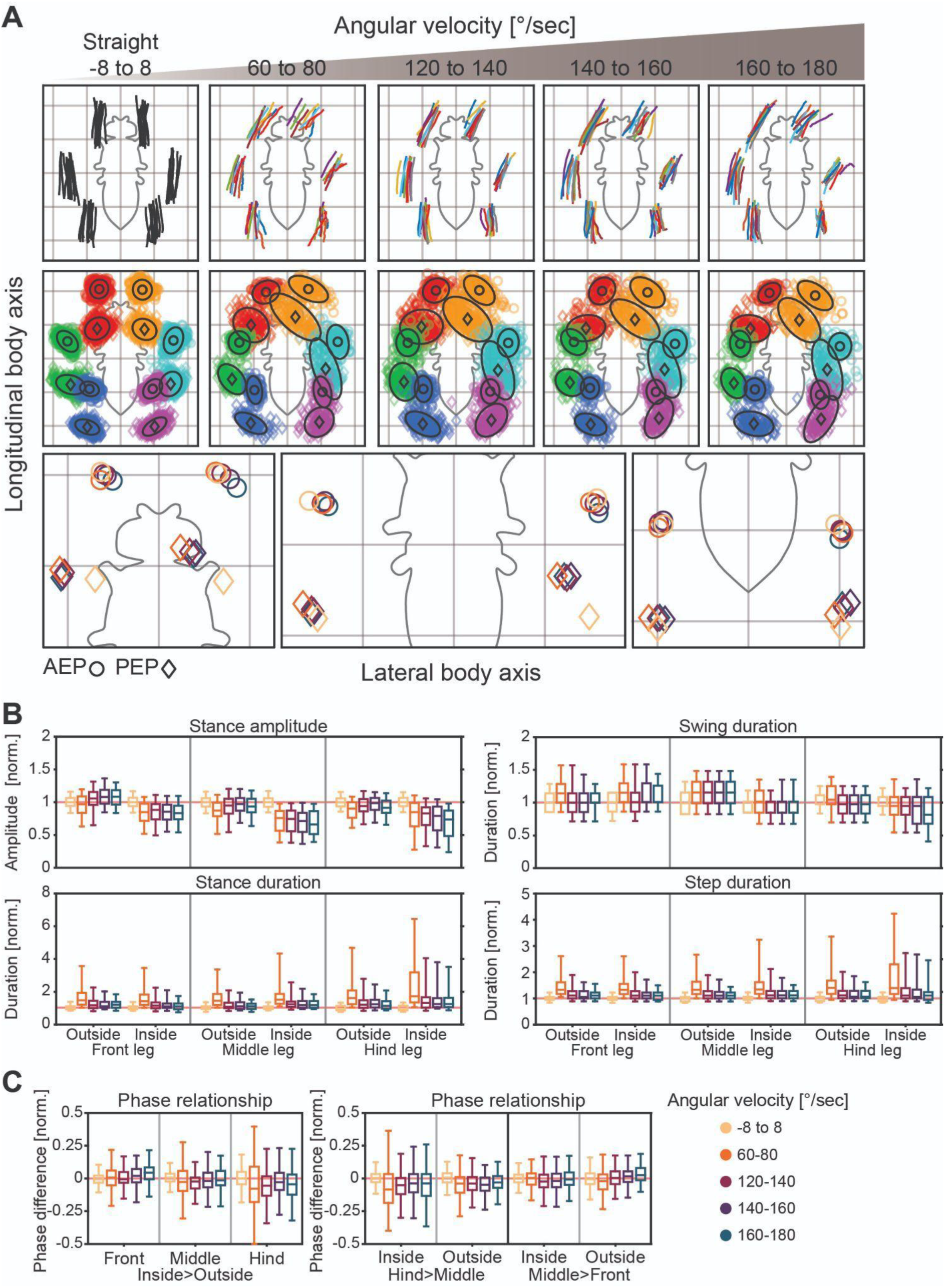
Low kinematic analysis of turns at different angular velocities. **A)** Average stance trajectories (top) and distribution of AEP (circles) and PEP (diamonds) of all flies walking straight (-8° to 8°, N=11, n=120), or performing turns (N=13, n=453). **B)** Kinematic parameters calculated for complete steps for straight walking, and turns. All data shown here have been normalized to the mean values in the straight walking condition (indicated as 1, light red reference line). Box plots (whiskers indicate 1% and 99% percentiles) are combined normalized data for all flies. **C)** Phase relationships between ipsilateral (Hind>Middle and Middle>Front) and contralateral leg pairs (Front, Middle, Hind, Inside>Outside), Inside and Outside, refers to the legs on the inside or outside of the curvature trajectory. Only a selection of the total range of explored angular velocities is shown here, see supplementary figure 3 and 4 for more details. N and n, are from the complete experiment.

During turns, PEP positions of all legs at the lowest turning velocities already matched the most extreme positions observed during arcs, and mostly showed minimal changes through subsequent increments of angular velocities, leaving velocity-dependent adjustments to occur predominantly via AEP shifts, particularly in the FL and ML (Fig. 3A, Fig. S2 middle). The OFL maintained its PEP at the lateral extreme while its AEP shifted toward the direction of the turn. Conversely, the IFL PEP remained proximal to the body, and subtly drifted inward-posteriorly, while its AEP displaced markedly in the same direction. The OML PEP retracted toward the straight-walking baseline as the turn sharpened while its AEP shifted inward-posteriorly, whereas the IML AEP shifted posteriorly accompanied by an inward PEP displacement. The HLs acted as a distinct exception to this AEP-driven modulation. While the OHL PEP subtly moved inward past the straight walking position as its AEP retained its position, the IHL underwent a unique, systematic contraction where both the AEP moved posteriorly and the PEP shifted inward-anteriorly, markedly shortening the step amplitude (Fig. 3A, Fig. S2 middle).

The previously described shifts in AEP and PEP directly manifested as a systematic, asymmetrical reconfiguration of stance amplitude as angular velocity increased. The OFL exhibited a progressive increase in amplitude, while the IFL displayed a corresponding decrease. The most pronounced adjustments occurred on the IML and IHL. The IML showed a sharp, linear reduction in amplitude, to approximately 50% of the straight-walking baseline at peak angular velocities, while the IHL similarly contracted its step to serve as a pivotal anchor (Fig 3B, Fig S4). In contrast, the OML and OHL maintained relatively stable amplitudes, probably providing the necessary forward propulsion. Amplitudes were calculated based on the distance between AEP and PEP; interestingly, however, we found that for the IML and IHL the stance trajectories were not always straight or slightly curved. In some cases, any or both legs described a stance trajectory with an L or V shape (see video 4 and Fig S2C Turning angle > 100°).

Temporal changes were characterized by leg-specific modulations of phase durations (Fig. 3B, Fig. S4). While swing durations of the OFL and OHL remained unaltered, the middle segment exhibited a velocity-independent asymmetry: the OML showed an increase in swing duration, whereas the IML was consistently shorter. The IHL displayed a velocity-dependent decrease in swing duration at high angular velocities (>140°s^−1^). These swing modulations were paired with substantial adjustments in stance duration. Across all legs, and most prominently in the inside HL, stance durations were initially lower than in straight walking (with median near 2-fold for all legs), likely partially reflecting lower translational speeds, but also skipped steps (for the IML and IHL), before progressively decreasing as angular velocity increased. However, at the highest angular velocities, this trend reversed specifically in the inside HL, where stance duration became higher once more. This temporal profile in the HL suggests a critical transition in motor strategy, potentially marking the threshold where the fly switches stepping strategies (further analyzed in the following sections, see Fig 6). Given the magnitude of stance duration changes, total step duration of each leg mainly reflected these changes

Phase relationships during turns exhibited segment-specific dynamics that both parallel and diverge from arc maneuvers (Fig. 3C, Fig. S4). In the front segment, the contralateral FL phase difference hovered tightly around zero across almost the entire velocity range. Conversely, and consistent with our observations during arcs, the contralateral ML showed a slight negative phase shift uncoupled from velocity increments, except at the highest angular velocity where the phase reversed. In contrast, the contralateral HL pair displayed a distinct, speed-independent decoupling profile; across the lowest angular velocities tested, the phase difference dropped below zero and subsequently fluctuated non-linearly across the velocity spectrum without a directed trend, remaining consistently negative while exhibiting a massive, persistent data dispersion.

Ipsilateral phase analysis during turns revealed a substantial expansion in data dispersion compared to arc maneuvers, a feature especially pronounced in the HL>ML inside pairing (Fig. 3C, Fig. S4). Both the HL>ML inside and outside couplings maintained phase differences predominantly below zero; the inside pairing exhibited a slight, velocity-dependent upward trend in its median. In the anterior segment, the ML>FL inside relationship remained near zero across the entire velocity spectrum, whereas the ML>FL outside phase difference displayed a progressive, velocity-dependent increase trend.

### Spatial scaling and temporal divergence in angle-dependent turns

While arcs are executed at near-constant velocities, making the cumulative turning angle a direct function of the number of steps, turns involve complex but short-lived dynamics, where the contribution of step number is tightly coupled with acceleration profiles and the duration of the high-velocity peak. Categorizing steering maneuvers by angular magnitude has been a traditional approach in insect locomotion. Early studies in honeybees (Zolotov et al. 1975), fruit flies (Strauss and Heisenberg 1990) and cockroaches (Jindrich and Full 1999) introduced angular thresholds to describe turning behavior, suggesting that sharper turns might rely on distinct motor strategies. To test whether heading change alone dictates leg-level coordination, and whether angle-defined maneuvers map onto the same kinematic control regimes identified in our dynamic analysis, we categorized our dataset by final heading angle and recalculated the same kinematic parameters.

Segregating sequences by turning angle revealed distinct leg-specific AEP and PEP displacement profiles (Fig. 4A, Fig. S2 bottom). The OFL AEP was generally located slightly posterior to the straight-walking baseline, whereas the IFL AEP subtly shifted inward for smaller angles and converged toward the baseline at larger angles. Meanwhile, FL PEP displacements mirrored the velocity-dependent trends seen in arcs, moving progressively outward with increasing turning angles. For the OML, the AEP shifted closer to the body without further changes, while the PEP moved slightly outward-anteriorly at larger angles. On the IML, the AEP hovered posterior to its baseline position, whereas the PEP initiated from an anterior, proximal position and retreated closer to the body as the heading angle scaled, echoing arc maneuvers. The HL also displayed axis-specific modulations. The OHL AEP and PEP remained practically unaltered, presenting minimal displacements without clear tendency, except for an anterior displacement of the PEP.. Conversely, the IHL AEP shifted non-linearly toward a posterior position, whereas the PEP exhibited a consistent anterior displacement, systematically closing the distance to the AEP as the final heading angle increased.

**Figure 4.**
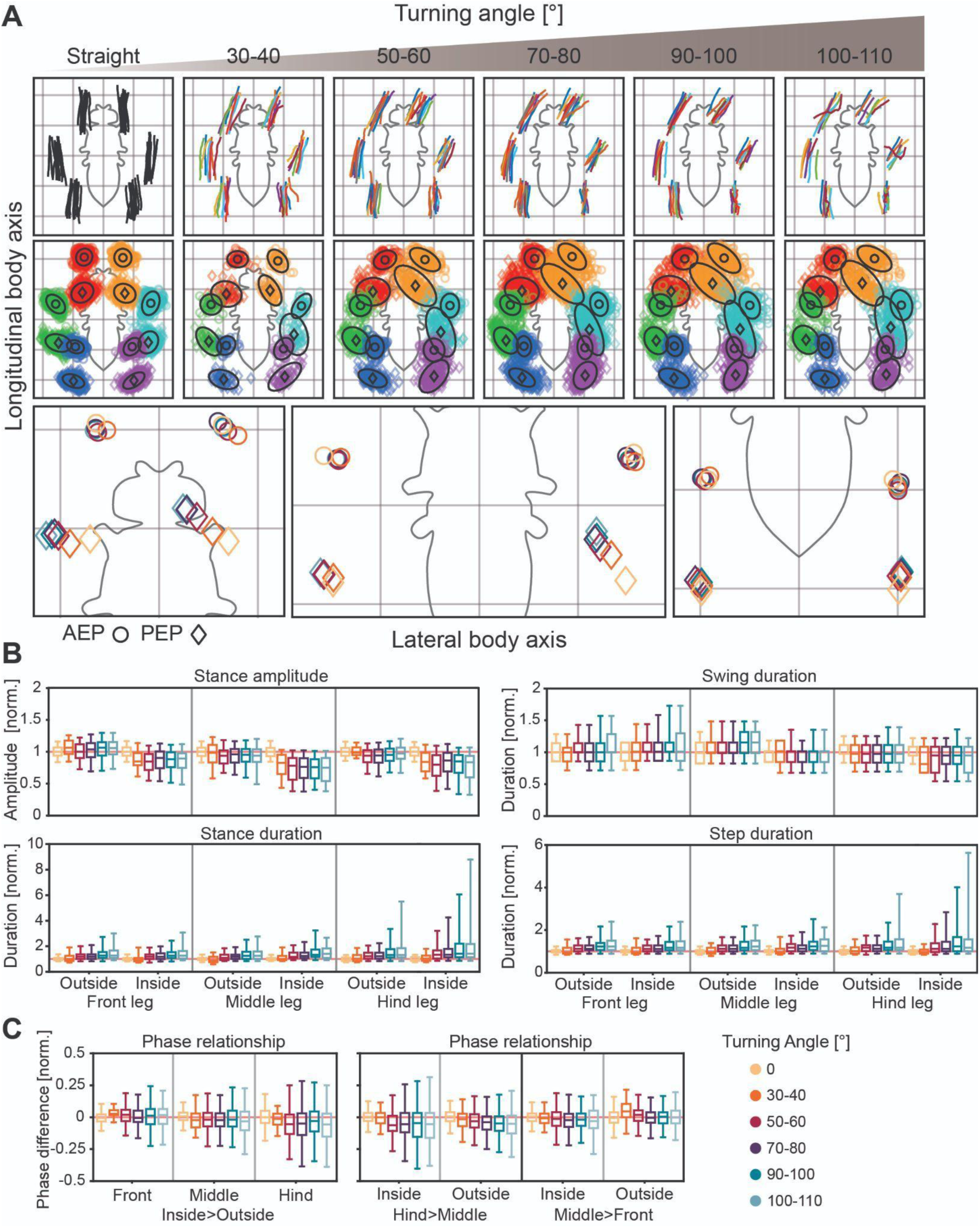
Low kinematic analysis of turns at different turning angles. **A)** Average stance trajectories (top) and distribution of AEP (circles) and PEP (diamonds) of all flies walking straight (0°, N=11, n=120), or performing turns (N=13, n=434). **B)** Kinematic parameters calculated for complete steps for straight walking, arc and turns. All data shown here have been normalized to the mean values in the straight walking condition (indicated as 1, light red reference line). Box plots (whiskers indicate 1% and 99% percentiles) are combined normalized data for all flies. **C)** Phase relationships between ipsilateral (Hind>Middle and Middle>Front) and contralateral leg pairs (Front, Middle, Hind, Inside>Outside), Inside and Outside, refers to the legs on the inside or outside of the curvature trajectory. Only a selection of the total range of explored turning angles is shown here, see supplementary figure 3 and 5 for more details. N and n, are from the complete experiment.

Ordering turning sequences by final turning angle revealed that stance amplitude and swing duration trends remained consistent with those observed across increasing angular velocities (Fig. 4B, Fig. S5). However, stance duration exhibits an inverse relationship to the trend seen before. As the final turning angle increased, the stance duration of all legs became progressively higher, with this increase being particularly pronounced in the HL (Fig 4B, Fig S5). This increase directly influenced the overall step duration, which increased accordingly. Regarding interleg coordination, the variability of the phase relationships increased continuously as a function of the final turning angle, largely maintaining the overall directional trend observed in the velocity-based analysis (Fig 4C, Fig S5).

In conclusion, comparing kinematic modulations driven by turning angle versus angular velocity reveals a fundamental divergence in temporal control. While both regimes share consistent trends in stance amplitude and swing duration, they deploy opposing temporal strategies. To achieve high angular velocities, *Drosophila* shortens the step cycle by drastically reducing stance duration. Conversely, to attain large net angular displacements, stance phase is extended, particularly in the hindlegs, prolonging step duration to sustain turning torque over time. Furthermore, the continuous accumulation of phase variability suggests that both high-speed and large-amplitude maneuvers represent highly flexible locomotor states where strict tripod coordination is progressively relaxed to accommodate mechanical constraints

### Backward-stepping turns engage multi-leg lateralization, extreme stance compression, and extended stance durations

We identified 70 turning sequences involving backward hindleg steps. Due to their specialized nature and lower relative abundance, these sequences were pooled for analysis, independent of angular velocity or final turning angle.

During these maneuvers, the IFL and IML executed lateral steps nearly perpendicular to the body’s longitudinal axis, with the IML mostly accompanying the IHL in backward stepping (Fig. 5A top, Fig. 6, and Video 5). Coupled with this backward movement, the amplitude of inside stance trajectories became progressively shorter; while the OFL trajectory amplitude remained unchanged relative to straight walking, all other trajectories shortened significantly, with inside segments displaying high variability and a massive data dispersion (Fig. 5B). Concurrently, the OFL executed more lateralized trajectories than in other turning categories, while the OML and OHL shifted along the direction of the curve and outward, respectively, collectively forming a circular path that reflected body rotation (Fig. 5A top). Across all legs, variance was highest in the PEP, especially for the inside legs in which backward steps occurred, causing inside AEP and PEP coordinates to shift closer together and capturing the extreme reduction in stance amplitude (Fig. 5A bottom)

**Figure 5.**
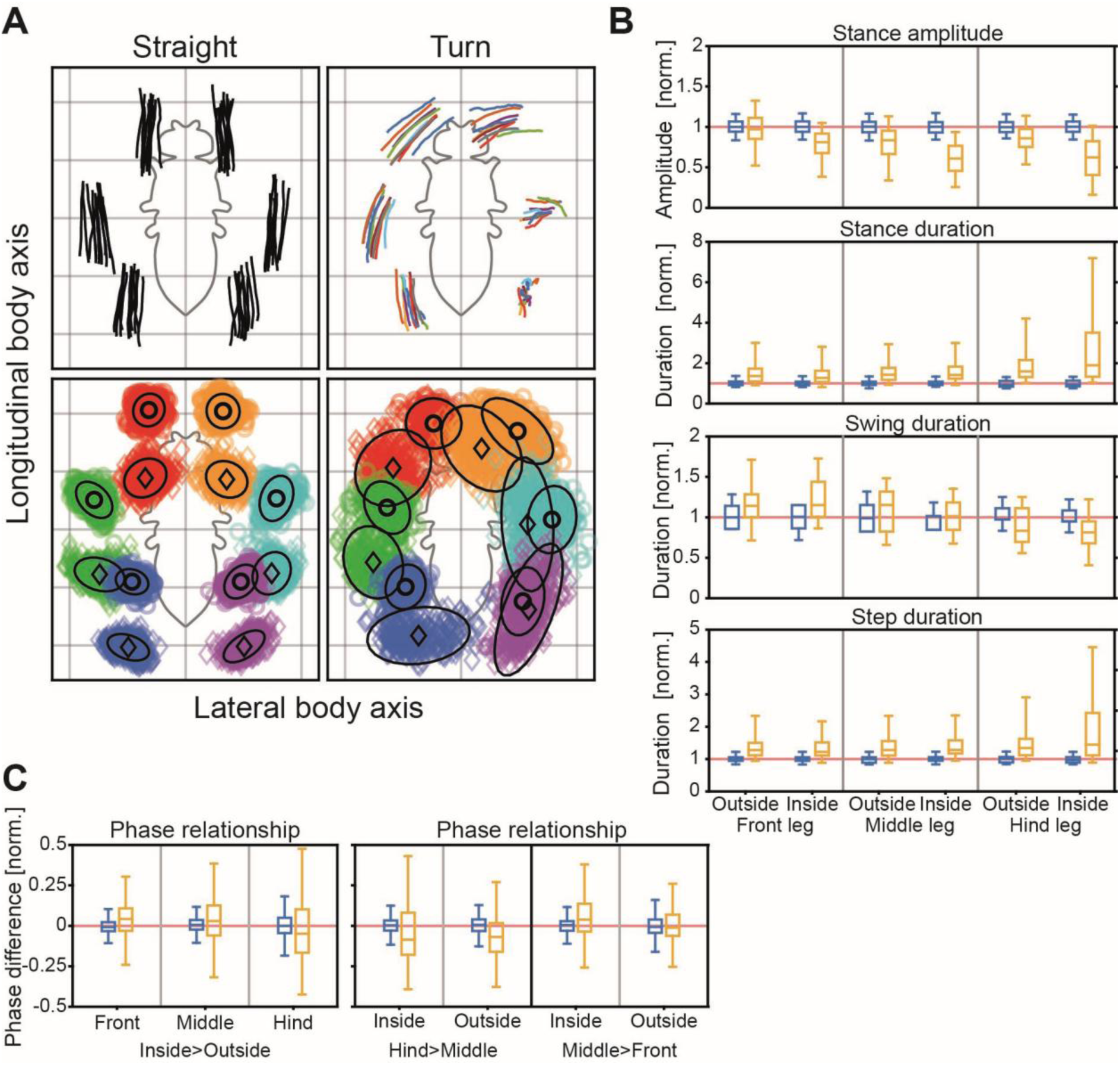
Low kinematic analysis of tight turns. **A)** Average stance trajectories (top) and distribution of AEP (circles) and PEP (diamonds) of all flies walking straight (N=11, n=120), or performing tight turns (N=11, n=70). **B)** Kinematic parameters calculated for complete steps for straight walking, arc and turns. All data shown here have been normalized to the mean values in the straight walking condition (indicated as 1, light red reference line). Box plots (whiskers indicate 1% and 99% percentiles) are combined normalized data for all flies. **C)** Phase relationships between ipsilateral (Hind>Middle and Middle>Front) and contralateral leg pairs (Front, Middle, Hind, Inside>Outside), Inside and Outside, refers to the legs on the inside or outside of the curvature trajectory.

**Figure 6.**
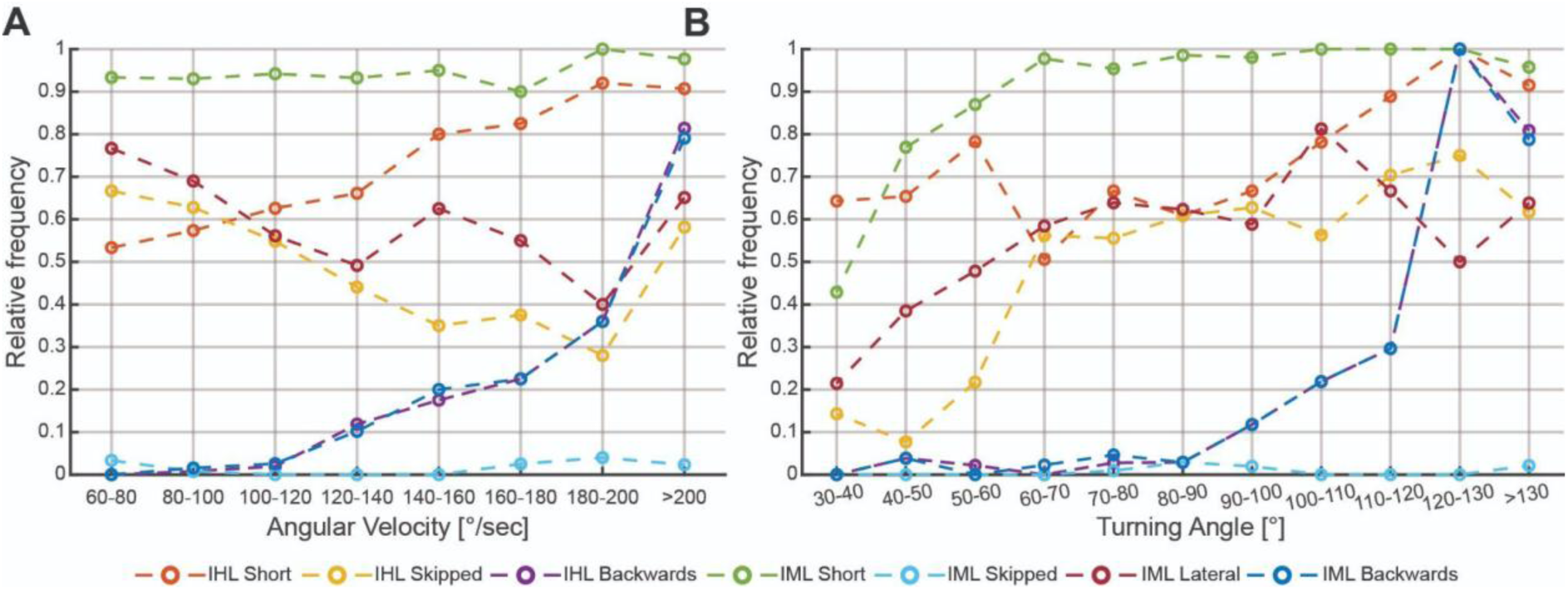
Relative frequencies of step strategies during turns at different A) Angular velocities, and B) Turning angles. IHL: Inside Hind Leg, IML: Inside Middle Leg. (N=13, n=513)

Swing duration variability increased across all segments. Both FL and the OML exceeded straight-walking durations, the IML remained unchanged, whereas the HL displayed an opposing trend, with both swing durations dropping below baseline and the IHL being the shortest (Fig. 5B). Overall step duration increased across all legs, driven by a dramatic extension of stance duration. This was most pronounced in the IHL, which occasionally reached seven times the baseline, likely reflecting multiple skipped steps.

While distinct directional shifts occur in median phase relationships, data exhibited a pronounced increase in overall variance, capturing a continuum of heterogeneous stepping behaviors (Fig. 5C, left). Between contralateral pairs, variability scaled progressively from FL to HL. Contralateral FL and ML pairs showed a positive shift above baseline, indicating a temporal phase advance of the outside leg, whereas the HL displayed an opposing negative shift, reflecting a delayed outside phase or a prolonged anchoring effect on the inside due to skipped steps. Ipsilateral coordination also underwent segment-specific modifications (Fig. 5C, right). Along both axes, the HL>ML phase relationship dropped significantly below zero, indicating a delayed phase transmission from posterior to middle segments. Conversely, the ML>FL phase relationship shifted above zero exclusively on the inside, capturing an accelerated phase coupling between the ML and FL on the inner side of the curve

### Hierarchical recruitment of segment-specific motor strategies during turning

Turning maneuvers encompassed a diverse repertoire of leg-specific motor adaptations beyond simple trajectory modulations. On the inner side of the curve, we identified seven distinct stepping strategies deployed by the ML and HL, including step shortening, step skipping, and backward steps, alongside lateral/perpendicular steps in the ML (Fig. 5A). To map how these local adjustments fulfilled higher-level behavioral requirements, we quantified the relative frequency of each strategy across systematic increments of both angular velocity and turning angle, pooling our standard turning dataset with sequences containing backward steps.

Analyzing locomotor strategy distributions as a function of angular velocity revealed a clear transition from localized kinematic modifications to a synchronized backward-stepping regime (Fig. 6A). At lower angular velocities (60 to 100°s^−1^), the behavioral output was highly dynamic. The relative frequency of IHL short-stepping rose steadily from ∼0.53, while IHL skipping decayed continuously from ∼0.67, culminating in a crossover near 100°s^−1^. Concurrently, the IML relied heavily on step shortening (∼0.93) while its lateral stepping initiated a progressive overall decline from ∼0.77, with backward stepping in both segments remaining virtually non-existent. As velocities increased to the intermediate range (100 to 180° s^−1^), backward steps in both the IHL and IML emerged simultaneously, rising from near zero to approximately ∼0.35. Throughout this same range, IML lateral stepping exhibited a progressive overall reduction punctuated by transient increases at 140 to 160° s^−1^ and above 200° s^−1^, respectively, reaching its minimum at 180 to 200° s^−1^. Beyond this 180 to 200° s^−1^ window, there was an abrupt, switch-like inflection: the frequency of backward steps in both the IML and IHL surged simultaneously from ∼0.37 to frequencies of ∼0.80. This demonstrates that at extreme angular velocities, *Drosophila* abandons standard forward adjustments in favor of a tightly coupled, multi-leg backward-stepping module.

A complementary organization unfolds according to the spatial magnitude of the maneuvers when evaluating strategy distributions as a function of turning angle (Fig. 6B). At lower turning angles (30° to 60°), the motor output transitions rapidly. IML short-stepping and lateral stepping rise sharply from ∼0.43 to ∼0.88 and from ∼0.22 to 0.48, respectively. Concurrently, IHL short-stepping increases from ∼0.64 to 0.78, while IHL skipping initiates a steep ascent from ∼0.15, with backward stepping remaining virtually absent.

Between 60° and 110°, relative frequencies level off. IML short-stepping saturates near its ceiling (∼0.95 to 1.0), while IML lateral stepping, IHL skipping, and IHL short-stepping stabilize around ∼0.60 to 0.70. At 100°, IHL short-stepping started increasing until reaching near 1 at 120°. Beyond 110°, the system undergoes a radical, synchronized inflection. The relative frequencies of backward steps in both the IML and IHL surge vertically from ∼0.30 to a peak near 1.0 at 120°-130°, rapidly dominating the motor output during the sharpest maneuvers.

## Discussion

### Arc and turn as distinct behavioral modes: separable categories within a graded kinematic space

Our identification of arcs and turns as two kinematically separable steering strategies in freely walking *Drosophila* aligns with previous descriptions in flies (Strauss and Heisenberg 1990; Yang et al. 2024) and other insects (Zolotov et al. 1975; Jindrich and Full 1999). Our quantitative separation of arcs and turns confirms that these maneuvers represent genuinely distinct behavioral modes. The finding that translational deceleration systematically precedes peak angular velocity during turns highlights a key mechanical constraint: flies, like other insects, must actively shed forward momentum to execute sharp-radius rotation without slipping (Jindrich and Full 1999). From a control perspective, this temporal dissociation suggests that steering is not controlled by a single descending drive, where a stronger signal would simply create tighter turns. Instead, it supports a dual-axis command architecture where translational speed and angular rotation are regulated via independent pathways, allowing the animal to decelerate the center of mass before a high-torque body rotation.

The relationship between arc kinematics and the established forward-walking locomotor manifold deserves explicit consideration. In straight walking, the locomotor manifold represents a low-dimensional, continuous geometric surface that constrains six-legged coordination, parameterized along its primary axis by stance duration scaling with translational speed, namely interleg coordination in *Drosophila* exists on a continuum largely characterized by speed-dependent changes in stance duration (Wosnitza et al. 2013; Szczecinski et al. 2018; DeAngelis et al. 2019). Because arcs were speed-matched to straight-walking controls, any kinematic differences reported here cannot be attributed to a speed shift along this continuum, but must reflect turning-specific aspects. Crucially, step duration remained constant across the full range of angular velocities in arcs (40 to 220°s^−1^), with swing and stance durations compensating for each other. This indicates that increasing curvature during arcs does not displace the animal along the speed-dependent axis of the locomotor manifold, meaning that executing a tighter arc does not require a change in forward speed. Instead, it recruits a spatially driven modulation, primarily PEP displacement and asymmetric stance amplitude, that operates independently of the temporal step cycle. This prioritization of spatial over temporal remodeling perfectly mirrors how *Drosophila* adapts to asymmetric speed perturbations on a split-belt treadmill, where flies corrected their course by shifting segment-specifically AEP and PEP positions, while stance duration remained unaltered (Pratt et al. 2024). Furthermore, the near-complete conservation of ipsilateral phase relationships suggests that the metachronal coordination rules governing straight walking remain intact during arcs (Wosnitza et al. 2013; Mendes et al. 2013; Cruse 1990). Together, these observations suggest that arc generation represents a feedforward spatial expansion of the forward-walking manifold into a curvature dimension orthogonal to speed, rather than a transition to a distinct locomotor state.

Although arc and turn trajectories are statistically separable, several parameters suggest an underlying spatial continuum spanning both behavioral categories. The stance amplitudes of the inside legs decrease monotonically with increasing rotational demand; this is true for velocity-sorted arcs and angle-sorted turns. When compared along the angular velocity axis, PEP positions at turn onset already match the most extreme values reached during arcs (Fig. 2, Fig. 3), indicating that the spatial boundary between strategies is not marked by a discrete jump in leg positioning, but by a shift in which each parameter (PEP or AEP) carries the primary steering signal. At the extremes, backward-stepping turns recruit nearly perpendicular IML steps (Fig. 5), representing the furthest point of this spatial progression. Because tarsal contact points during stance define the support polygon, the convex hull on the substrate that encloses the body’s center of mass, these leg-specific displacements directly reshape the animal’s stability geometry. Thus, the spatial reconfiguration of the support polygon is a graded process driven by geometric demand. However, while the transition from arcs to forward turns operating within the forward-stepping regime may represent a smooth extension of the forward manifold, we propose that this continuum has a strict functional boundary. The abrupt appearance of the multi-leg backward stepping module at high-turning demands, either high-speed or large-angle turns (Fig. 6), effectively represents a non-linear escape from the forward-walking manifold, dissolving PEP/AEP saturation constraints to transition the system into a distinct high-torque motor attractor.

### The PEP/AEP dissociation between arcs and turns reveals two distinct steering mechanisms

A key novelty of our findings is that *Drosophila* steering is achieved through two distinct phase-specific control mechanisms recruited hierarchically depending on turning demand. Importantly, they do not operate uniformly across all legs, but are implemented in a highly segment- and leg-specific manner. During gradual arcs, steering relies almost exclusively on a PEP-driven stance bias (mechanism 1). As steering demands become higher for sharp turns, the first mechanism saturates, and a AEP-driven swing redirection (mechanism 2) is sequentially recruited and superimposed on the mechanism 1.

During shallow-curvature steering (arcs), the motor system relies on a spatially graded modulation of the PEP. This mechanism alters the kinematics of stance trajectories, while the tarsus remains in ground contact, biasing the subsequent swing phase without reorganizing the core metachronal rhythm. This stance-phase tuning likely relies on local, proprioceptive feedback loops, such as load-sensitive campaniform sensilla (Zill et al. 2004; Duysens et al. 2000). Because step duration remains at straight-walking baseline levels (Fig. 2, Fig. S3) and the metachronal coordination rules remain intact, this PEP-driven mechanism represents a continuous spatial expansion of the forward-walking manifold rather than a transition to a distinct locomotor state.

In contrast, sharp turns require the recruitment of the second mechanism, to modify the stride through progressive AEP displacement. OncePEP modulation reaches immediate saturation at the transition to turns, further rotational torque is generated by altering AEP placement, directing the leg in the air to target a new touchdown position. This active redirection of swing-phase targeting likely involves, feedforward descending modulation of the gain and/or the sign of the proprioceptive feedback, that normally helps limit anterior swing/protraction and promote the swing-to-stance transition. This could include feedback from hair plates as well as chordotonal organs (Mamiya et al. 2018; Ulrich Bässler 1988; Büschges 1989; Hofmann et al. 1985; Hofmann and Koch 1985; U. Bässler 1993; Kuenzi and Burrows 1995; Wong and Pearson 1976; Dean and Schmitz 1992; Pratt et al. 2026). During these maneuvers, the outside limbs decouple from the rotational geometry to focus on propulsion, while the inside ML and HLcontract sharply to pivot the body (Fig. 4).

Notably, the transition to backward-stepping turns drives the IML to PEP and AEP coordinates far beyond the forward-stepping turn limits (Fig. 5). This demonstrates that the PEP saturation observed at the arc-to-turn boundary is not an anatomical constraint, but rather a functional boundary of the forward-walking manifold. By actively recruiting a synchronized, multi-leg backward-stepping module, the nervous system dissolves this boundary. This transition migrates the functional pivot point directly onto the insect’s body axis, enabling near-in-place rotation. Whether this boundary is dictated by physical stability, propulsive limits, or descending command thresholds remains to be elucidated. Thus, whereas forward steering is systematically parsed into a PEP-driven stance bias and an AEP-driven swing redirection, the activation of the backward-stepping module represents a distinct motor program that overrides both phase-specific regimes.

### A dual-axis, hierarchical framework links descending steering commands to segment-specific kinematics

Our analysis reveals a fundamental temporal trade-off in HL kinematics: while maximizing angular velocity requires reduction in stance duration, achieving large turning angles relies on an increase. This behavioral dissociation between angular velocity and turning angle in our dataset cannot be predicted by a simple, linear rate-code model of steering, where a single descending command of increasing frequency proportionally scales both rotational speed and angle, as classically modeled in bilateral descending drive (Yang et al. 2024). Instead, it exposes two independent temporal axes of motor control operating primarily at the level of hindleg stance duration, which adapt the temporal structure of the step depending on whether the animal optimizes for the speed of rotation or the completeness of an orientation maneuver. Recent circuit-level analyses in Drosophila provide a possible neural substrate for this dual-axis control: the descending neuron DNa03 acts as an effector-agnostic integration hub, transforming navigational cues into an abstract velocity signal that anticipates maneuvers across both walking and flight (Feng et al. 2026). For walking, this signal is integrated over time by the downstream node DNa11, effectively encoding the accumulation of heading change at the top of the walking execution layer to reorient all six legs during high-amplitude turns. Finally, fine-grained walking kinematics are implemented by subordinate descending neurons, with DNa02 driving inside stride shortening and DNg13 driving outside stride lengthening, translating independent velocity and heading commands into coordinated execution (Yang et al. 2024).

The opposing motor behaviors in HLs observed during angular velocity or heading optimization directly mirror the distinct temporal profiles of DNa03 and DNa11. Under the first axis of high instantaneous rotational speed, high DNa03 drive actively reduces IHL stance duration to increase stepping frequency, maximizing the rate of turning torque generation per unit of time. Conversely, under the second axis of cumulative heading reorientation, sustained integration by DNa11 increases IHL stance duration, prolonging ground contact and sometimes inducing skipped steps to stabilize the body’s turning pivot while the contralateral legs sweep the body around. Thus, the temporal axis of motor control, stance duration, in the anchoring HL is differentially and opposingly modulated by descending networks depending on which navigational demand is prioritized.

This dual-axis temporal control is closely coupled with a segmented and hierarchical motor organization where spatial parameters are modulated in a highly leg- and segment-specific manner. During gradual steering, like in arcs, the system operates within a continuous, forward-walking regime where these two axes remain largely independent. Here, turning is achieved through subtle, phase-specific spatial asymmetries, primarily driven by PEP adjustments in the IML and IFL, altering the stance trajectories, while keeping the metachronal rhythm and step durations identical to straight-walking controls (Fig 2B, S2A, S3). This low-level course correction is likely implemented via low-order descending pathways like DNa02 and DNg13 (Yang et al. 2024), allowing the insect to drift smoothly without reorganizing its core interleg coordination. However, as steering demand increases and these spatial PEP adjustments saturate, the system transitions to a more “aggressive” regime dominated by AEP repositioning (Fig 3; S2B).

Under this sharp turning regime, the primary control of steering shifts to the swing phase, actively redirecting the inside legs to target a more posterior touchdown position, while the OML and OHL partially decouple from the turning geometry to focus purely on forward propulsion (Fig. 3, Fig. 4, S2B-C). At the limits of turning demand, where standard forward kinematics can no longer satisfy the reorientation requirements, the system undergoes an abrupt, non-linear switch to a synchronized, multi-leg backward-stepping module (Fig. 6). Kinematically, this maneuver engages a coordinated multi-leg lateralization: while the IHL transitions to backward stepping, both the IFL and IML execute lateral steps nearly perpendicular to the body’s longitudinal axis, with the IML additionally accompanying the IHL in active retraction and backward stepping (Fig. 5). This combined spatial reconfiguration shifts the turning pivot directly onto the fly’s longitudinal axis, enabling near-in-place rotation. Our finding that the ML is recruited actively during backward stepping is strongly supported by connectomic and physiological evidence showing that DNa11 commands individual thoracic segments in parallel rather than relying on mechanical substrate coupling (Feng et al. 2026). Biomechanically, this sudden transition represents an active, segment-specific reflex reversal functionally analogous to the central-complex-mediated transitions characterized in larger insects (Martin et al. 2015). Under strong descending DNa11 influence, load-related and joint-position feedback that normally promote forward propulsion are dynamically re-routed at the local premotor level to drive active retraction of the inside limbs (Ruthe et al. 2026; Akay et al. 2007; Gruhn et al. 2016).

Remarkably, these independent pathways and hierarchical layers appear to operate primarily under conditions of intermediate turning demand, where they can flex the path of the fly independently during natural navigation, such as odor-plume tracking. However, at maximum angular velocities and turning angles from our dataset, the distribution of individual stepping strategies and their associated temporal parameters converge almost identically (Fig. 6). This phenotypic alignment could indicate that at extreme demands, the dual control axes co-saturate downstream motor networks, driving the system into a single, saturated motor state. Regardless of whether the initial drive originates from instantaneous velocity demands (DNa03) or cumulative heading requirements (DNa11), maximal recruitment of the steering hierarchy forces the downstream motor system into a single, highly stereotypical kinematic bottleneck. Under the uniform optomotor stimuli utilized in laboratory assays, these commands are driven to their co-saturating limits, exposing the fixed-point attractor states that define the hard mechanical and neurobiological boundaries of hexapod maneuverability.

In conclusion, integrating velocity and angle distributions reveals that *Drosophila* turning behavior is governed by a hierarchical, dual-mode control strategy. While shallow steering maneuvers rely on proportional, segment-specific adjustments of forward kinematics, extreme spatio-temporal turning manoeuvres trigger a non-linear, switch-like transition characterized by the synchronous activation of a multi-leg backward-stepping module that overrides standard coordination. The functional divergence of IHL skipped steps demonstrates that while static anchoring is prioritized to sustain geometric wide-angle turns, it is actively dismissed in favor of dynamic backward propulsion when shifting toward high rotational speeds. Remarkably, comparing the endpoints of both analyses reveals a striking convergence in stepping frequencies at the limits of locomotor performance (Fig. 6A, >200°s^−1^ vs. Fig. 6B, >130°). At these extreme operational boundaries, individual strategy frequencies align almost identically across both panels.This precise mirroring demonstrates that the insect’s motor control network recruits a shared, stereotypic steering configuration during maximum behavioral demands, suggesting that distinct spatial and temporal steering pathways ultimately drive the motor system toward a common kinematic output.

## Resource availability

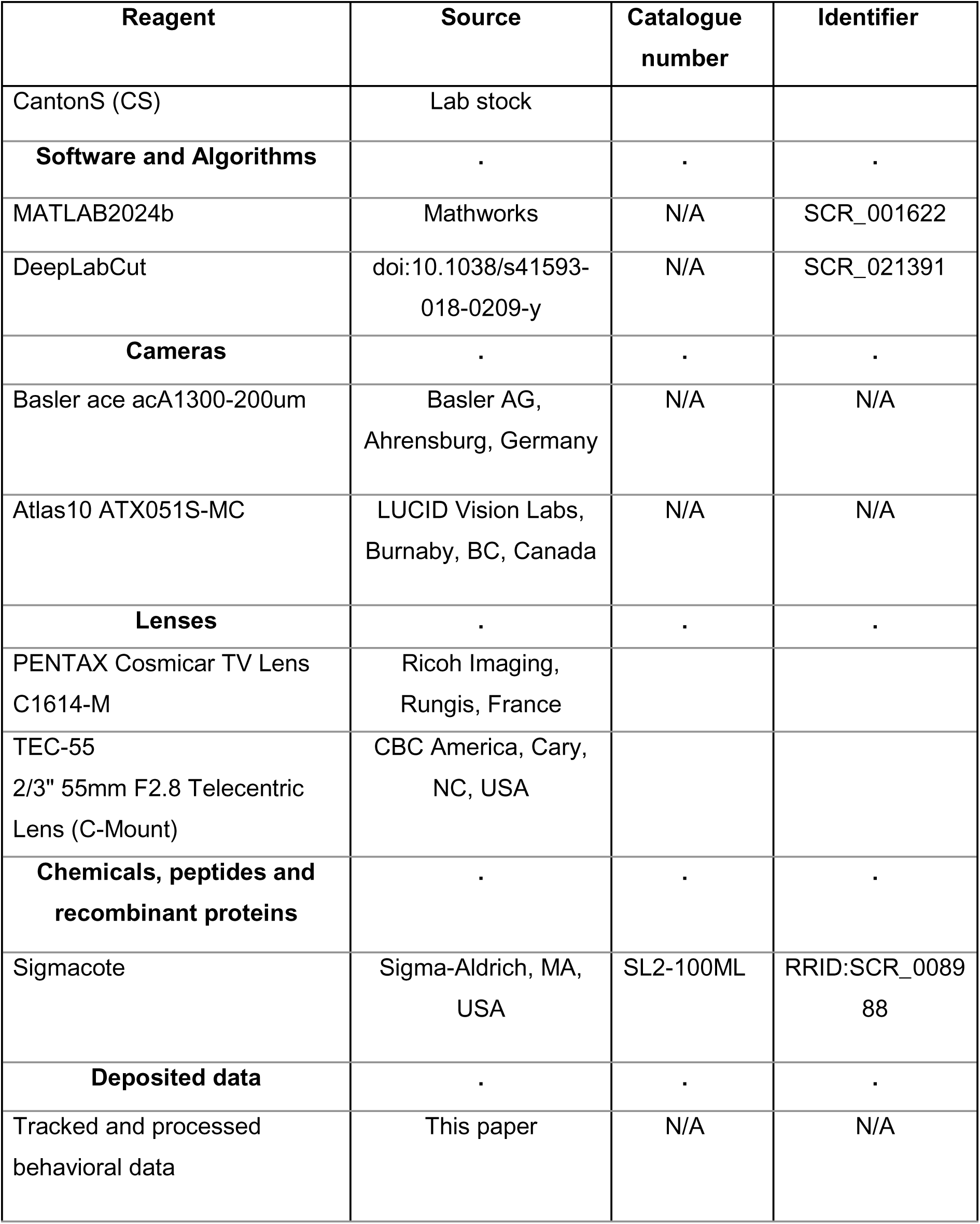

## Experimental model and study participant details

### Fly stock

*Drosophila melanogaster* female flies aged 3-5 days were used for this study. Flies were reared on a standard medium at 25°C on a 12 h:12 h light:dark cycle, and 50-70% humidity.

All experiments were carried out on *Canton S* flies.

## Method details

### Experimental setups

The experimental setups were custom designed and built by our Electronics workshop, Department of Animal Physiology, University of Cologne. Behavioral data collection was done using two similar setups with technical differences.

### Behavioral setups

Two versions of the same setup were employed to capture fly behavior. The first version was described in depth in previous studies (Chockley et al. 2022; Godesberg et al. 2024), and was used only for figure one. The final output of these setups is undistinguishable, and can be analysed with the same tools. The second version differs from the previous one mainly on the use of an Atlas ATX051S-MC camera (LUCID Vision Labs, Richmond, BC, Canada), also equipped with an object-space telecentric lens (focal length 55 mm, model: TEC-55, CBC America, Cary, North Carolina, USA). Due to its two times higher resolution and the 10 Gigabit Ethernet connection, this camera allowed us to continuously grab frames with a resolution of 2000-by-2000 pixels at a recording rate of 200 Hz, capturing the whole arena continuously. The walking arena consisted of an inverted glass Petri dish, of 60mm diameter with a watch glass of 100mm as the lid. The inner side of the watch glass was coated with SigmaCote (SL2, Sigma-Aldrich, St Louis, MO, USA) to reduce the ability of the flies to walk on it. This petri dish was placed in a custom-made plastic holder. The camera was located on the side of the setup and it was directed at the experimental chamber from below using a mirror tilted at 45 deg. The setup was placed in dim ambient light, mostly dark. The chamber was illuminated with a ring of 60 infrared (IR) LEDs (wavelength 890 nm, opening angle 20 deg). To eliminate any remaining ambient light from the recordings, and enhance the contrast, a visible-light filter (upper cut-off frequency 790 nm) was added to the camera’s lens. Acquisition of individual video frames during this shutter time and IR illumination were synchronized with a pulse generator.

To overcome the bottleneck created by fast storing and handling the massive amount of data generated, we wrote and implemented a new Python software. This software recognizes the fly while capturing the video, and only keeps the part of the image in which it is located. Briefly, it first downsamples video frames by a factor of 4, then it converts them into binary images with a brightness threshold, and then it applies a median function to the white pixel values along the x- and y-axes. Afterwards, the center of the largest object with brightness values above threshold (the fly) can be detected in less than 5 ms and with an accuracy of 4 pixels. Around the position of the fly a 200-by-200 pixel square is cut out of the video frame. On this new section, the position of the fly is again determined, but without downsampling to have more exact coordinates and a smoother walking trajectory. The resulting 200-by-200 pixel cut is then written to the hard drive. In addition to it, the exact position of the section in the original video frame was stored to keep track of the walking trajectory and to enable reconstruction of the original view. Exact time points of recording for each frame were saved to enable the post hoc detection of frame drops. In this case, a BL of around 2.5mm corresponds to 105 pixels.

#### Processing of recorded behavior and extraction of sequences of interest

The resulting data sets from the new setup contained a broad set of behavior that a fly can produce in it, ranging from grooming to walking. An additional Python software was developed in order to algorithmically filter these data sets. Three parameters were defined: (1) walking speed, (2) curvature of the walking trajectory, and (3) distance to the center. All three parameters were calculated and assigned to each individual video frame. In addition, a minimum trajectory length was established to filter out segments that, while valid, were too short for analysis purposes.

1. Walking speed was calculated by measuring the distances between the fly positions every 21 frames to smooth out speed fluctuations which happen when legs transition between swing and stance phase. An additional benefit of smoothing was the reduction of noise caused by both the discretization inherent in the detection algorithm and the brightness fluctuations of the moving fly body. These fluctuations can affect which pixels surpass the brightness threshold, thereby influencing the determination of the fly position.
2. The curvature was approximated by fitting a spline function to an equidistantly resampled version of the trajectory, followed by evaluating this spline to compute a curvature value at each point along the original trajectory. The application of the spline function effectively filtered out minor bumps, wiggling of the fly body, and artifacts from the detection algorithm, while preserving genuine changes in the direction of movement. To acquire a basis for the spline function to work on, the original walking trajectories were rearranged into equidistant pieces. This means that the original trajectory was resampled and after every 0.25 mm a new anchor point was taken. These anchor points then formed the equidistant trajectory and were used to calculate separate spline functions for the x- and y-components. These separate spline functions could be combined afterwards, resulting in a smoothed version of the original trajectory, which left out smaller bumps and spikes in the trajectory but reflected changes in the overall direction of movement. Because this smoothed trajectory was fully based on the spline functions, these functions could directly be used to calculate the curvature at any point of the trajectory.
3. For the calculation of the distance to the center the position of the arena in the original camera frame needed to be estimated. For this purpose, the smallest circle containing all recorded fly positions was calculated and the center of that circle was the approximation of the center of the arena. The distance of each point on the walking trajectory to this approximated arena center was calculated to enable subtraction of every behavior which happened close to the rim of the arena.

We adjusted these parameters to extract from each original video, sequences of straight walking, arcs, and turns. After extraction, sequences were visually inspected to avoid false classification.

Then, videos from both setup versions were used for annotation of eight different body parts in every video frame: the tarsal tips of all six legs, the neck and the end of the abdomen. This was automatically performed in DeepLabCut (DLC; (Mathis et al. 2018)), and inspected afterwards to correct mis-annotations.

### Behavioral experiments

To avoid anaesthesia effects on behavior a single fly was transferred into the arena, by aspirating it in a tube and gently letting it walk into the chamber. Flies were then allowed to acclimate to the setup for 5 min prior start video acquisition, which duration depended on the experimental condition. During this time, flies walked free and spontaneously in the chamber.

### Optomotor stimulation assay for silencing experiments

Surrounding and centered on the FlySpector setup a modular LED display was placed (adapted from (Reiser and Dickinson 2008)). It formed a cylindrical arena composed of 100 8×8 Blue (Φ3.0mm, 470nm) LED panels (KWM-30881ABB/ KWM-30881CBB, Luckylight), arranged in a 5×20 grid (Reiser and Dickinson 2008). Each square panel has 32.0 mm height, forming a cylindrical display of 16 cm height and approximately 20 cm diameter around the setup. The LED displays were controlled using the MATLAB script PControl (https://reiserlab.github.io/Modular-LED-Display/) (Reiser and Dickinson 2008). A pattern consisting of alternating 5 LED wide (15mm on the display) vertical bars the full height of the display was presented during the experiments. Intensity of the bars was set on level 2 of the PControl options (0 to 7 levels). Each experiment consisted of 5 phases, 30min each. First, a phase where bars were presented as a stationary pattern to an individual fly, from which control straight walking sequences were collected. Afterwards, the pattern started to rotate counterclockwise at 4 different speeds, 10, 20, 30, and 50 LEDs displacement per second (3, 6, 9, and 15 cm/s on the screen). This was done to increase the probability of the fly performing arcs and turns with different features. Between phases in which the bars were rotating, we included a 5 min resting phase, in which bars were steady. Extraction and selection of the videos were done as explained before. Angular velocity and turning angle were calculated after DLC annotation and correction.

### Free-walking kinematic analysis

To determine the times of lift-off and touch-down for each leg, the DLC-determined positions of the tarsal tips were transformed into a world-centered coordinate system. In this coordinate system, when the leg is on the ground during a stance phase its speed is close to 0 mm/s, while during a swing phase it moves with respect to the ground. These measures and an empirically determined threshold were used to distinguish swing and stance movements. Transitions between these two phases were defined as touch-down and lift-off events, in which tarsal tips positions were defined as the anterior and posterior extreme positions (AEP and PEP) in the fly-centered coordinates. For each leg a step was defined as its movement between two consecutive PEPs; its period was defined as the time difference between them. Swing phase and its duration were defined as the movement and the time difference between a PEP and the following AEP; stance phase and its duration were defined as the movement and the time difference between an AEP and the following PEP. A stance trajectory was defined as the complete path of a tarsal tip in fly-centered coordinates between an AEP and the following PEP. Step amplitude was defined as the distance between a PEP and the following AEP. Average AEPs and PEPs were defined as the arithmetic mean of all AEP and PEP position vectors, respectively; the standard deviation of these positions was estimated as a bivariate distribution.

Individual shifts in extreme positions during directional changes were calculated as vectors between the mean AEPs/PEPs in the control condition and their respective individual AEPs/PEPs during the directional change; these can be described with anatomical directions (posterior, anterior, medial and lateral). Stance trajectories were averaged by first resampling all *n* trajectories to 100 equidistant 2D positions (100×2 data points) and then calculating the arithmetic mean for each set of 100×2×*n* data points. To simplify the analysis and the interpretation of the results all AEP and PEP positions were flipped when required, so that all curve-walking segments used in the analyses were performed to the same side (left in this paper).

### Free-walking phase analysis

Phase relationships (i.e. phase differences) were calculated for all contralaterally and ipsilaterally adjacent leg pairs. For contralateral phase relationships the left leg was chosen as the reference leg; and for ipsilateral phase relationships, it was the posterior leg of the pair. For each complete step of the reference leg we calculated its instantaneous phase as a value that linearly increased from 0 to 1 during the step. We then calculated the phase relationship between the two legs as the phase value at the times of PEPs in the non-reference leg; this was equivalent to the phase difference between these legs. A value of 0 indicated synchronous lift off, while a value of 0.5 indicated an anti-phase relationship. The phase between two legs is given as, for instance, Hind>Middle, where Hind is the reference leg and Middle is the non-reference leg.

Aside from DLC-based functions (for details, see (Mathis et al. 2018)), all annotations, calculations and statistics for the kinematic as well as the phase analysis were carried out with custom-written functions in MATLAB 2024b (RRID:SCR_001622; see also ‘Statistical analyses’, below) and the MATLAB CircStat toolbox (Berens 2009).

### Angular velocity

angular velocity was calculated as the average change in body orientation for each frame of walking trajectory. The calculation of body orientation was based on determining the main body axis, defined by neck and abdomen markers, annotated using DLC. The difference in body orientation between each frame of each trajectory was measured and the resulting value was presented in °s-1.

### Turning Angle

First we determined the maximum value of curvature in a turning trajectory, and its position within the trajectory. Afterwards a parametric linear fit was performed to the segments of the trajectory before and after the maximum curvature value, and the fit was vectorized to properly assess the angle of change in direction. Finally, the angle created between the two vectors was calculated.

### Statistical analyses

To facilitate comparison between the straight walking and turning conditions, we normalized all individual step periods, swing durations, stance durations and stance amplitudes to the arithmetic mean of the straight condition. Flies differ naturally with regard to these parameters, and normalization allows for direct comparison between flies and pooling of all data. It should be noted that, because of the extensive dataset and the correspondingly high sample size, many of the tested differences are highly significant.

#### Receiver Operating Characteristic and high kinematic parameters

To assess the discriminability of Arc and Turn walking strategies at the individual trial level, we performed a Receiver Operating Characteristic (ROC) analysis across nine kinematic parameters: mean translational speed, distance traveled, number of steps, mean angular velocity, standard deviation of curvature, standard deviation of cumulative heading, maximum cumulative heading deviation, angular velocity per unit distance, and angular velocity variance. For each parameter, its value was used as a continuous classifier score to discriminate Arc from Turn trials. ROC curves were computed using MATLAB’s *perfcurve* function, sweeping all possible classification thresholds. When the resulting Area Under the Curve (AUC) was below 0.5, the curve was reflected, yielding a direction-agnostic measure of discrimination. AUC values were interpreted as follows: ≥0.9 excellent, ≥0.8 good, ≥0.7 fair, ≥0.6 poor, and <0.6 no discrimination above chance. Group differences for each parameter were additionally assessed using the Mann-Whitney U test, with effect sizes quantified as the rank-biserial correlation coefficient r = 1 − 2U/(n₁n₂), where |r| thresholds of 0.1, 0.3, and 0.5 corresponded to small, medium, and large effects respectively. All analyses were performed in MATLAB (MathWorks). Kernel density estimates of parameter distributions were computed using MATLAB’s *ksdensity* function with default bandwidth selection for all three strategies (Arc, Turn, and Straight). For parameters where the Straight strategy exhibited an interquartile range below 5% of the total IQR across all groups, indicating near-zero variance relative to the overall data range, Straight was excluded from the corresponding distribution plots to avoid axis compression that would obscure biologically relevant variation in Arc and Turn trajectories.

## Acknowledgments

We thank Mehrdad Ghanbari, Sima Seyed-Nejadi and Sherylane Seeliger for technical assistance, and the members of the Büschges Lab for their input on the project.

## Funding

AB and KI were supported by research grant C3NS, part of the Next Generation Networks for Neuroscience Program (DFG grants Bu857/15-1,2; NSF DBI grant 436258345), by DFG grant CRC1451 (SFB1451/1,2, 431549029) and by ‘iBehave’ funded by the Ministry of Culture and Science of the State of North Rhine-Westphalia. KI was supported by the AXA Chair of Research Fund.

## Author contributions

- EAG, AB, and TB conceived and designed the research.
- EAG, MD, and TB developed the behavioral setups.
- EAG and RDC performed experiments and analyzed data.
- TB developed scripts and software for behavioral recordings and data analysis.
- EAG, RDC, TB, and AB interpreted the results.
- EAG, TB, and AB designed and prepared figures.
- EAG drafted the manuscript.
- EAG, AB, and TB edited and revised the manuscript. All authors approved the final version of the manuscript.
- KI and AB provided resources and funding.

## Declaration of interests

The authors declare no competing interests.

---

**Supplementary figure 1.**
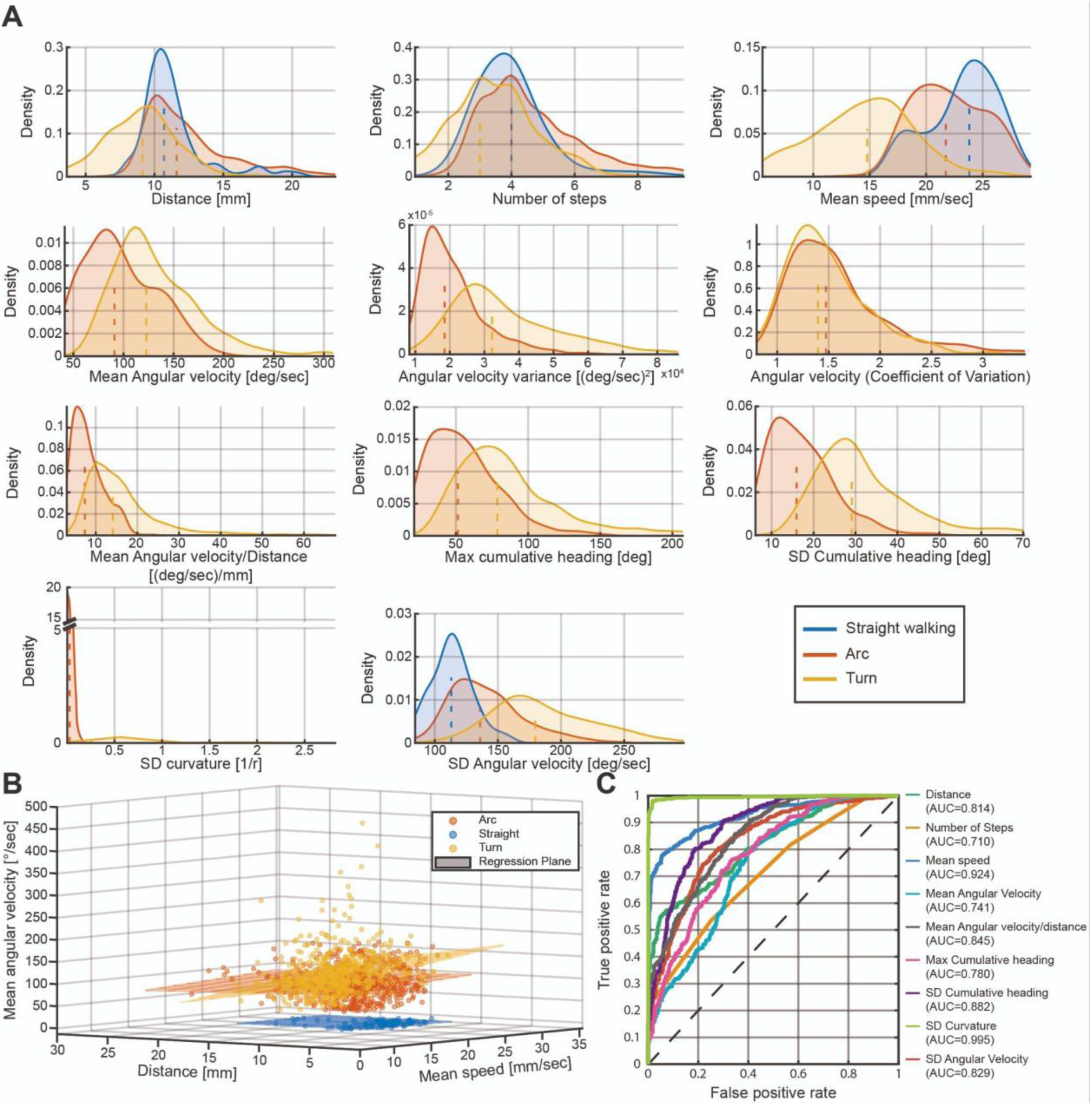
**A)** Kernel Density Curves of several parameters of straight, arcs and turns sequences. Straight was excluded from plots when its interquartile range was below 5% of the total data range (max-min across all groups) for that parameter. Straight: N= 11, n= 118; Arcs: N= 12, n= 527; Turns: N= 13, n= 522 **B)** Average angular velocity, distance, and average translational speed during the sequences. **C)** Receiver Operating Characteristic (ROC) of the different parameters evaluated. AUC = Area Under the Curve.

**Supplementary figure 2.**
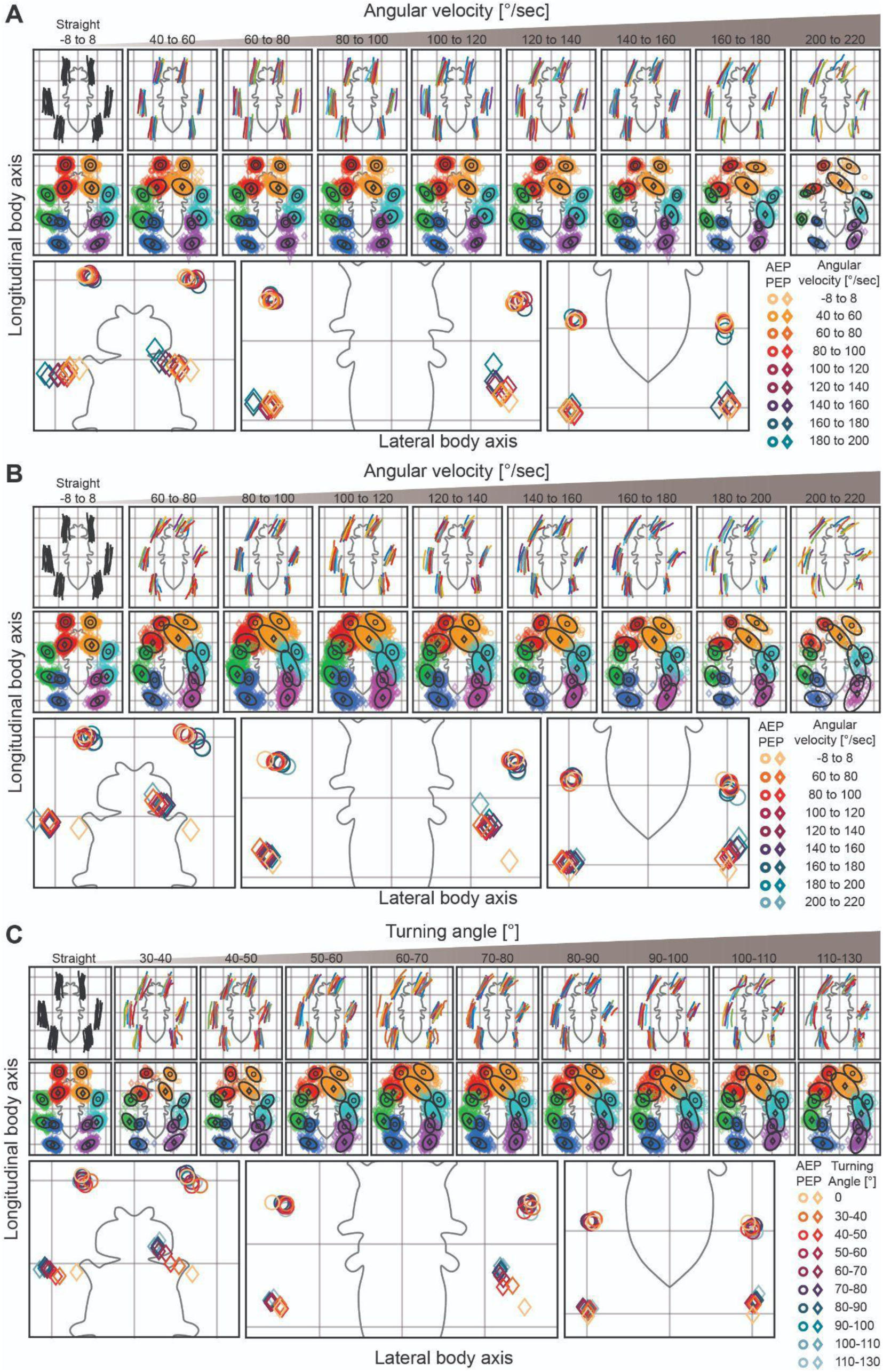
Average stance trajectories (top) and distribution of AEP (circles) and PEP (diamonds) in a fly-centered coordinate system of all flies walking straight (-8° to 8°, N=11, n=120), or performing **A)** arcs (N=11, n=527), **B-C)** turns (B: N=13, n=458; C: N=13, n= 434). Walking sequences were grouped according to **A-B)** Angular velocities, **C)** Turning angles.

**Supplementary figure 3.**
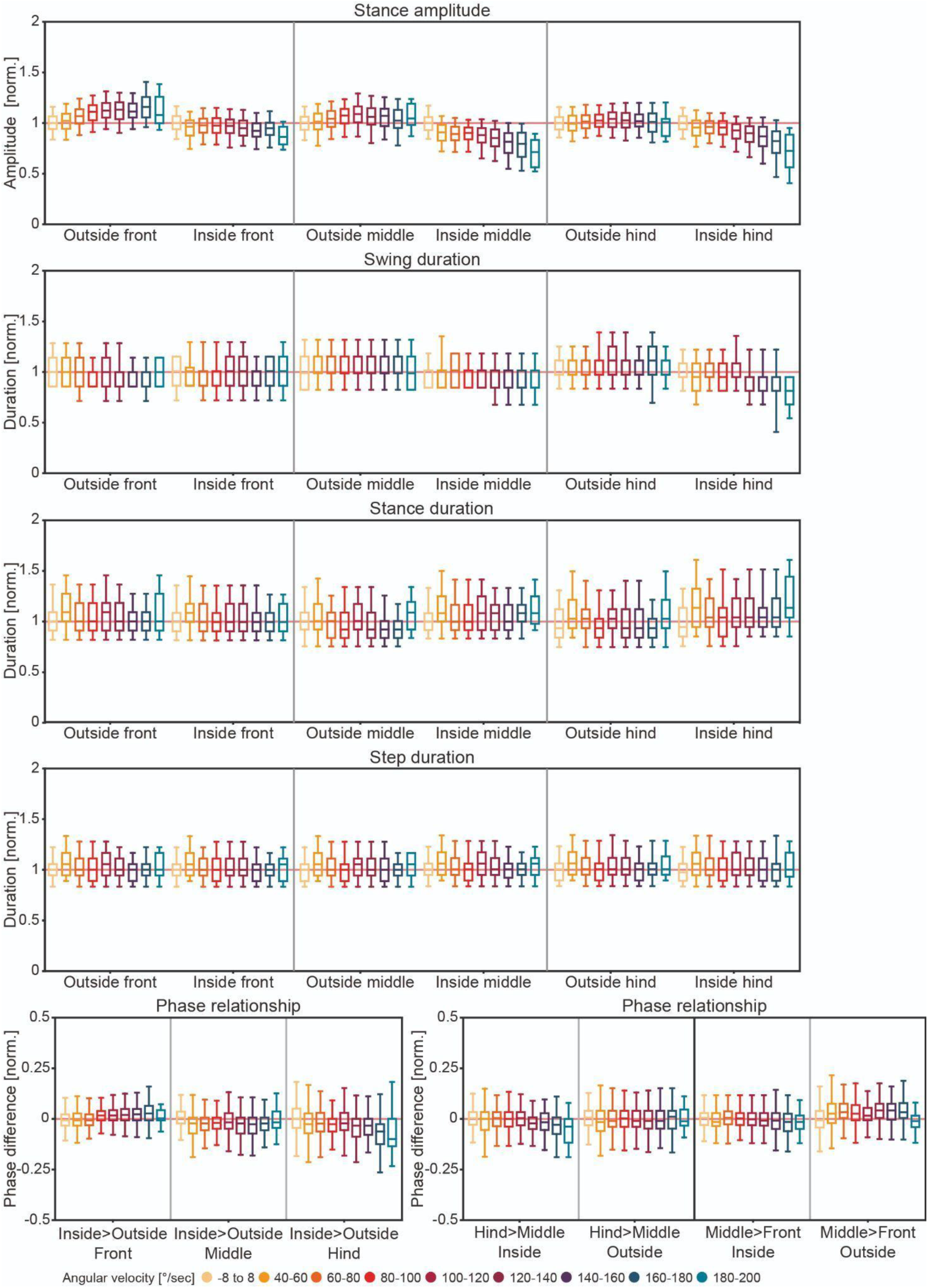
Low kinematic analysis of arcs at different angular velocities. Kinematic parameters calculated for complete steps for straight walking (-8° to 8°, N=11, n=120), and performing arcs (N=11, n=527). All data shown here have been normalized to the mean values in the straight walking condition (indicated as 1, light red reference line). Box plots are combined normalized data for all flies (whiskers indicate 1% and 99% percentiles).

**Supplementary figure 4.**
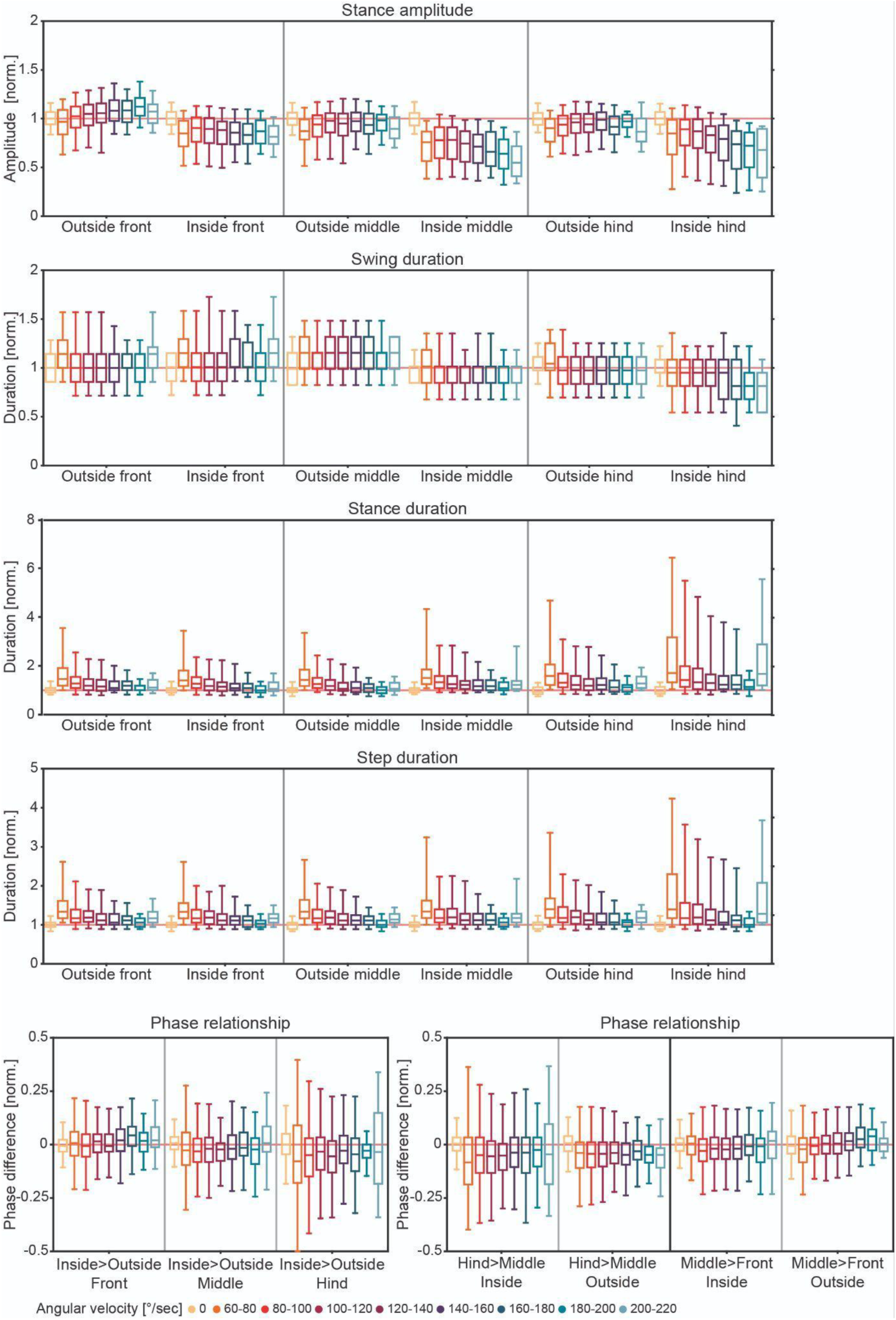
Low kinematic analysis of turns at different angular velocities. Kinematic parameters calculated for complete steps for straight walking (-8° to 8°, N=11, n=120), and turns (N=13, n=458), organized according to their angular velocity. All data shown here have been normalized to the mean values in the straight walking condition (indicated as 1, light red reference line). Box plots are combined normalized data for all flies (whiskers indicate 1% and 99% percentiles).

**Supplementary figure 5.**
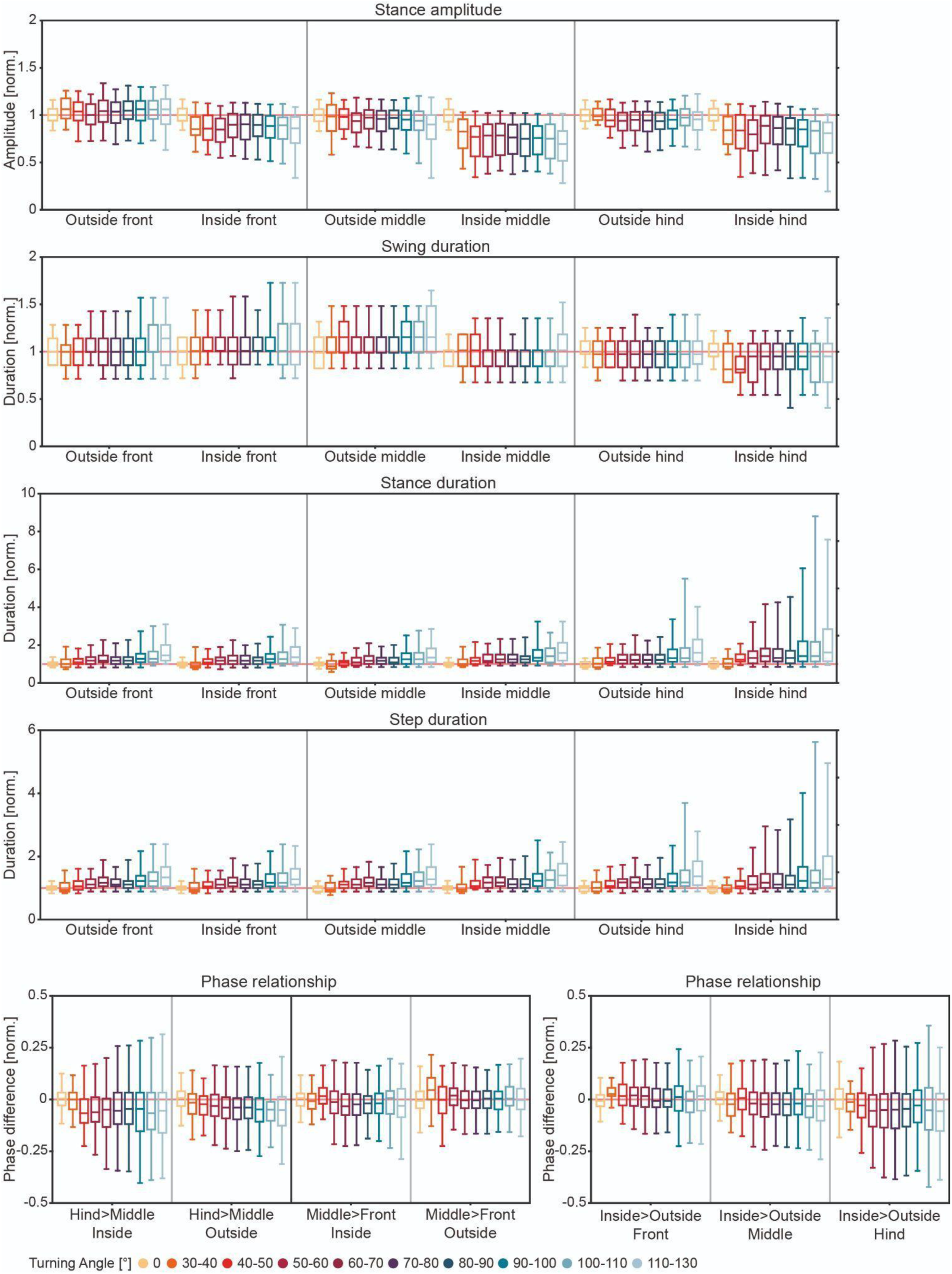
Low kinematic analysis of turns at different turning angles. Kinematic parameters calculated for complete steps for straight walking (-8° to 8°, N=11, n=120), and turns (N=13, n=434). All data shown here have been normalized to the mean values in the straight walking condition (indicated as 1, light red reference line). Box plots are combined normalized data for all flies (whiskers indicate 1% and 99% percentiles).

## Notes

### Competing Interest Statement

The authors have declared no competing interest.

